# Quantifying the Limits of Immunotherapies in Mice and Men

**DOI:** 10.1101/759167

**Authors:** Liam V Brown, Eamonn A Gaffney, Ann Ager, Jonathan Wagg, Mark C Coles

## Abstract

**Background:** CART-cells have demonstrated clinical success for the treatment of multiple different lymphomas and leukaemias. However, comparable clinical efficacy has not been observed for various solid tumours in human clinical trials, despite promising data from murine cancer models. Lower effective CART-cell delivery rates to human solid tumours compared to haematological malignancies in humans and solid tumours in mice might partially explain these divergent clinical outcomes, requiring the development of quantitative methods to calculate effective dosing in the clinic and models.

**Methods:** We used anatomical and physiological data for human and rodent circulatory systems to calculate the typical value and expected variation of the perfusion of healthy and tumour tissues, and used these to estimate the upper limits of immune cell delivery rates (maximum possible under optimal conditions) across different organs, tumours types and species.

**Results:** Estimated maximum delivery rates were up to order 10,000-fold greater in mice than humans yet reported CART-cell doses are typically only 10-100-fold lower in mice, suggesting that the effective delivery rates of CART-cells into tumours in clinical trials are far lower than in corresponding mouse models. Estimated delivery rates were compared to published PET data and found to be consistent.

**Conclusions:** The results suggest that higher effective human doses may be needed to drive efficacy comparable to mouse solid tumour models. These increased doses raise safety and manufacturing concerns. We posit that quantitation of species and organ-specific delivery and homing of engineered T-cells will be key to unlocking their potential for solid tumours.

## 1 Introduction

Cellular therapies such as CAR (Chimeric Antigen Receptor) T-cells have shown clinical efficacy against several leukaemias and lymphomas [1, 2]. This success has not yet been matched for solid tumours, despite the efficacy seen in pre-clinical models, and a suitable dosing strategy to maximise efficacy remains uncertain [3–8]. Typical response curves (amount of CART-cell transgene observed in blood versus time) in patients with haematological disorders are marked by an initial cellular expansion (typically 100-1000-fold [9]), due to the large numbers of CART and target cells colocalising in readily accessible tissues. Expansion increases the effective cellular dose entering and proliferating within compartments with lower perfusion or less efficient access, which can drive the clearance of target cells required to achieve complete responses in these compartments. In solid tumours, relatively few target cells are in readily accessible compartments, whether due to poor perfusion or barriers to extravasation, preventing a strong initial expansion of CART-cells. Tumour regression is achieved when the rate of tumour clearance is greater than that of tumour growth, including in the least perfused/accessible tumour lesions. In this context, tumour clearance is a numbers game and the relative lack of success for solid tumours may be due to lower effective CART-cell doses, since the number of accessible target cells is too low to drive the early cellular expansion that, in the case of haematological malignancies, increases the effective dose.

The delivery rate of cells to different compartments of the body will likely be of importance in CART-cell or eTCR (engineered T-cell receptor) responses. For intravenous (iv) administration, cells are delivered by the circulatory system. Systematic quantitation of the variation of vascular delivery rates across organs, tumour types and species will improve understanding of comparative preclinical and clinical outcomes and inform improved dosing strategies. Physiologically-based pharmacokinetic modelling (PBPK) has been used extensively to predict drug concentration profiles and their variability across different tissues and individuals, to estimate the efficacy of clinical dosing regimens (for recent reviews, see [10–12]). PBPK models have also been used in drug development since 2000 and are readily accepted as providing supporting information by both the US Food and Drug Administration and the European Medicines Agency. They have been further implemented in the investigation of T-cell trafficking, for example to determine the strength of the abscopal affect and influence of metastases on the primary tumour [13, 14] and to study localisation of adoptively transferred T-cells or cellular therapies [15–20]. However, we have not seen such models be used for quantitative exploration of the simpler consequences of differences between anatomical parameters in different species, nor an attempt to quantify and compare the maximum likely values of delivery rates of immune cells across organs and species, the aim of the present work.

We have made simple comparisons of the human, mouse and rat circulatory systems, using relevant organ, tumour and anatomical data [21–26]. We have calculated the upper bounds of cellular delivery from the circulation into each organ, considering only tissue perfusion and not factors that subsequently reduce rates of T-cell entry, such as tissue-specific extravasation probabilities or inflammation. The validity of predictions was tested through comparison to PET and imaging data taken shortly after cellular transfer, and the validity of maximum delivery rates for tumour tissue was found by comparing the typical perfusion of tumour and normal tissues. Predicted maximum delivery rates exhibited extreme differences by species. The delivery rate of cells per minute per mm^3^ to lungs is 20,000-fold higher in mice than humans, yet typical doses of CART cells given to experimental mice are only 100-fold less than those in the clinic. This may partially explain the lack of success seen against solid tumours reported to date.

## 2 Methods

### 2.1 Model summary

#### Model

Most studies of physiologically-based pharmacokinetics (PBPK) or cellular kinetics (PBCK) make use of an ordinary differential equation (ODE) model representing the anatomy. A schematic of the anatomy appropriate for such equations is shown in Figure 1. T-cells are assumed to flow from the heart to the vasculature of different organs, where they then return or extravasate into that organ’s interstitial space. Extravasated cells return to circulation via the lymphatics, except for the spleen and the pulmonary circuit, from which cells return directly. To calculate *maximum* delivery rates, we consider the case in which cells extravasate only in the tissue of interest, always extravasate once delivered by the vasculature, and do not return. This maximum rate is simply the rate at which cells are delivered by the vasculature, *i.e.* perfusion (blood flow ***B*** over total organ volume 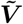) multiplied by blood concentration *C*. More precisely,

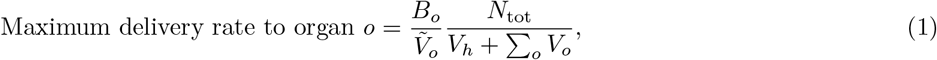

where *N*_tot_ is the total number of cells of interest, *V_o_* and 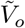 are the vascular and total volumes of organ *o*, and *V_h_* is the volume in the heart and interconnecting blood vessels. This expression can be shown to be equivalent to a special case of standard PBPK/PBCK models, see supplementary sections A.1 and A.1.4. This simplified scenario is presented in the inset of Figure 1. One organ is defined as the tumour bearing organ, containing a 1mm^3^ tumour (‘tmr’) tissue volume. We consider this volume either as healthy or tumour tissue, to find how predicted delivery rates to each differ across organs and species.

**Figure 1:**
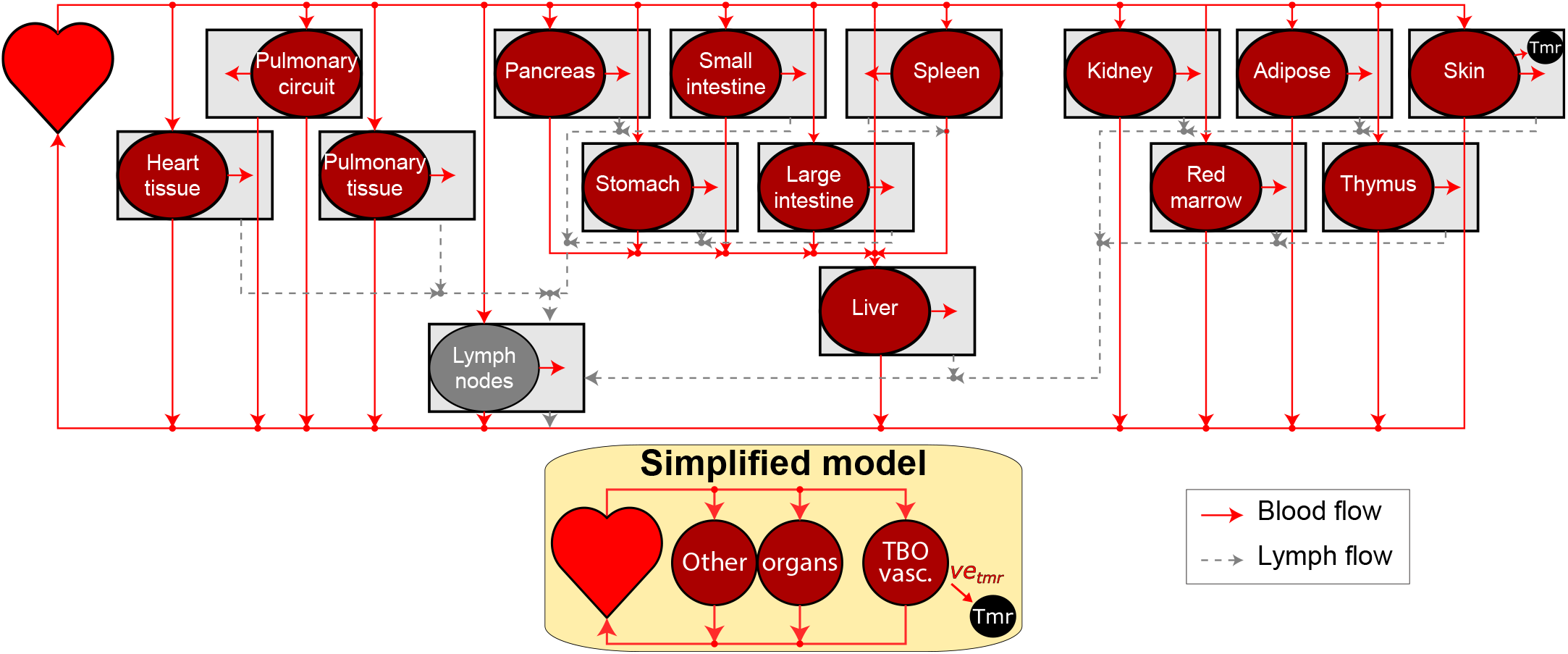
Model schematic. A visual summary of a model of the circulatory system. Solid and dotted lines represent blood and lymph flow respectively. Cells flow from the heart to each organ *o*, from which a proportion *e_o_* enters the interstitial space. Cells from the interstitium flow via the lymphatics back to the heart. One organ is designated as the tumour bearing organ (TBO), from which a proportion of cells *νe*_tmr_ extravasate irreversibly to the tumour (“Tmr”) instead of the organ interstitium, where 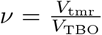 is the proportion of the TBO volume taken up by the tumour. The skin is set to be the tumour bearing organ as an example. **Inset, below:** When calculating the maximum rate of extravasation to a tumour by organ, a simplified version of the model that ignores much of the interstitial space was used, in which cells of interest are assumed not to extravasate to tissues other than the tumour. Details such as the portal vein are still present in the simplified model but not shown, for clarity.

### 2.2 Parameter selection from literature

Predicted T-cell delivery rates are dependent on assumed anatomical parameters (blood flow, blood volume and organ volume). We collected several anatomical reference banks from the literature [21–26], in particular the compilations by the ICRP and Shah *et al* [24, 26]. Each source has slightly differing fractional blood flows and volumes. To remove selection bias, delivery rates were calculated with many random values of anatomical parameters (*n* = 100 per organ per species), selected uniformly from the range of literature values, after which the means and standard deviations of estimated delivery rates were taken. This also serves as a proxy for population variability. To avoid using data from different studies for a single model animal, data sets that are as complete as possible were chosen. In particular, the total blood flow and blood volume, the volume of each organ, and the fractional blood flow and blood volume of each organ were recorded from each reference. These data are shown in supplementary tables S2, S3 and S4. Presented results are the mean and standard deviation of predictions obtained by choosing random values from the literature. Random parameter values are selected from the range of literature values. We cannot be more confident in any one report than another, so we choose the random values for all parameters (for each organs and species) uniformly. This process is repeated 100 times to yield the presented results. When considering tumour perfusion distinct from healthy organ perfusion, we use measurements of tumour perfusion from the literature (see supplementary table S3) and suppose that, since these are all measurements of different tumours, the data should follow a normal distribution. Thus, we choose normally distributed random values of tumour perfusion.

### 2.3 Generation of presented results

Presented data are maximum delivery rates in each species for each organ *o*, calculated using Equation 1, with some deviation due to details of the vasculature. For example, the portal vein blood flow must be added to *B_o_* for the liver (see supplementary section A.1 for further information). The results of Table 1 are obtained by applying data reported by Shah *et al* [24] to Equation 1. The results of Table 2 are obtained by multiplying the ratio of mouse to human delivery rates by the dose administered to mice, 10^7^.

**Table 1:**
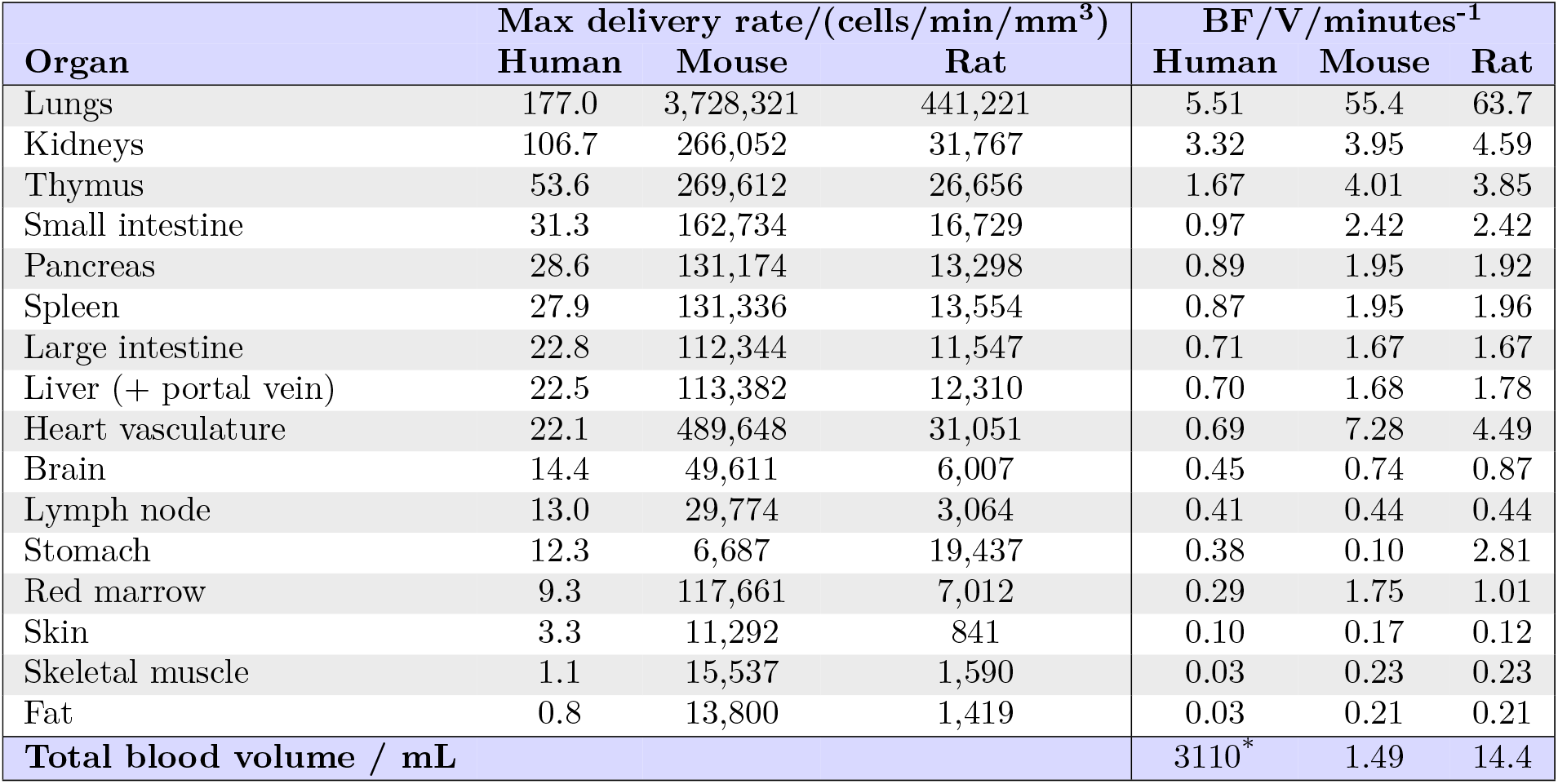
**Left: predicted absolute maximum CART-cell delivery rates per volume**(in cells/min/mm^3^) to non-tumour tissue in organs in humans, mice and rats, using previously compiled physiological parameter values [24]. It is assumed that organ perfusion is homogenous and 10^8^ CART-cells are introduced to each species. The interspecies differences in absolute delivery rates per volume depend only on organ perfusion and cell blood concentration. **Right: organ perfusion** (blood flow / organ volume; BF/V) and the total blood volume in each species, obtained by summing relevant volume data from [24]. *Note that the total blood volume from this reference is an underestimate, but it is expected to be underestimated by a similar amount in each species. The left table can be generated from the right by the formula 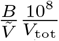, where *B* and 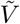 are the organ blood flow and volume and *V*_tot_ is the total blood volume in each species; see section 2.1.

| Organ | Max delivery rate/(cells/min/mm<sup>3</sup>) |  |  | BF/V/minutes<sup>-1</sup> |  |  |
| --- | --- | --- | --- | --- | --- | --- |
|  | Human | Mouse | Rat | Human | Mouse | Rat |
| Lungs | 177.0 | 3,728,321 | 441,221 | 5.51 | 55.4 | 63.7 |
| Kidneys | 106.7 | 266,052 | 31,767 | 3.32 | 3.95 | 4.59 |
| Thymus | 53.6 | 269,612 | 26,656 | 1.67 | 4.01 | 3.85 |
| Small intestine | 31.3 | 162,734 | 16,729 | 0.97 | 2.42 | 2.42 |
| Pancreas | 28.6 | 131,174 | 13,298 | 0.89 | 1.95 | 1.92 |
| Spleen | 27.9 | 131,336 | 13,554 | 0.87 | 1.95 | 1.96 |
| Large intestine | 22.8 | 112,344 | 11,547 | 0.71 | 1.67 | 1.67 |
| Liver (+ portal vein) | 22.5 | 113,382 | 12,310 | 0.70 | 1.68 | 1.78 |
| Heart vasculature | 22.1 | 489,648 | 31,051 | 0.69 | 7.28 | 4.49 |
| Brain | 14.4 | 49,611 | 6,007 | 0.45 | 0.74 | 0.87 |
| Lymph node | 13.0 | 29,774 | 3,064 | 0.41 | 0.44 | 0.44 |
| Stomach | 12.3 | 6,687 | 19,437 | 0.38 | 0.10 | 2.81 |
| Red marrow | 9.3 | 117,661 | 7,012 | 0.29 | 1.75 | 1.01 |
| Skin | 3.3 | 11,292 | 841 | 0.10 | 0.17 | 0.12 |
| Skeletal muscle | 1.1 | 15,537 | 1,590 | 0.03 | 0.23 | 0.23 |
| Fat | 0.8 | 13,800 | 1,419 | 0.03 | 0.21 | 0.21 |
| Total blood volume / mL |  |  |  | 3110* | 1.49 | 14.4 |

**Table 2:** **Human-equivalent dosages** for delivery to non-tumour tissue: The dosage of CART-cells in humans predicted to be required to give the same absolute delivery rate per mm^3^ as in a mouse given 10^7^ cells. The numbers required are much larger than many clinical dosages [27, 28].

| Organ | Equivalent dose | (continued) |  |
| --- | --- | --- | --- |
| Lungs | $1.7 \times 10^{11}$ | Heart vasculature | $2.2 \times 10^{11}$ |
| Kidneys | $2.5 \times 10^{10}$ | Brain | $3.4 \times 10^{10}$ |
| Thymus | $5.0 \times 10^{10}$ | Lymph node | $2.3 \times 10^{10}$ |
| Small intestine | $5.2 \times 10^{10}$ | Stomach | $5.4 \times 10^{09}$ |
| Pancreas | $4.6 \times 10^{10}$ | Red marrow | $1.3 \times 10^{11}$ |
| Spleen | $4.7 \times 10^{10}$ | Skin | $3.4 \times 10^{10}$ |
| Large intestine | $4.9 \times 10^{10}$ | Skeletal muscle | $1.4 \times 10^{11}$ |
| Liver | $5.0 \times 10^{10}$ | Fat | $1.7 \times 10^{11}$ |

Random results in Figure 2 are obtained by drawing uniformly random values of organ parameters (*n* = 100), calculating the maximum delivery rate with Equation 1, and subsequently finding the mean and standard deviation of delivery rates. *n* = 100 values were chosen for each organ to generate an indication of delivery rate variability, whilst ensuring that the mean of selected random parameters was within 5% of the actual mean of experimental parameter values.

**Figure 2:**
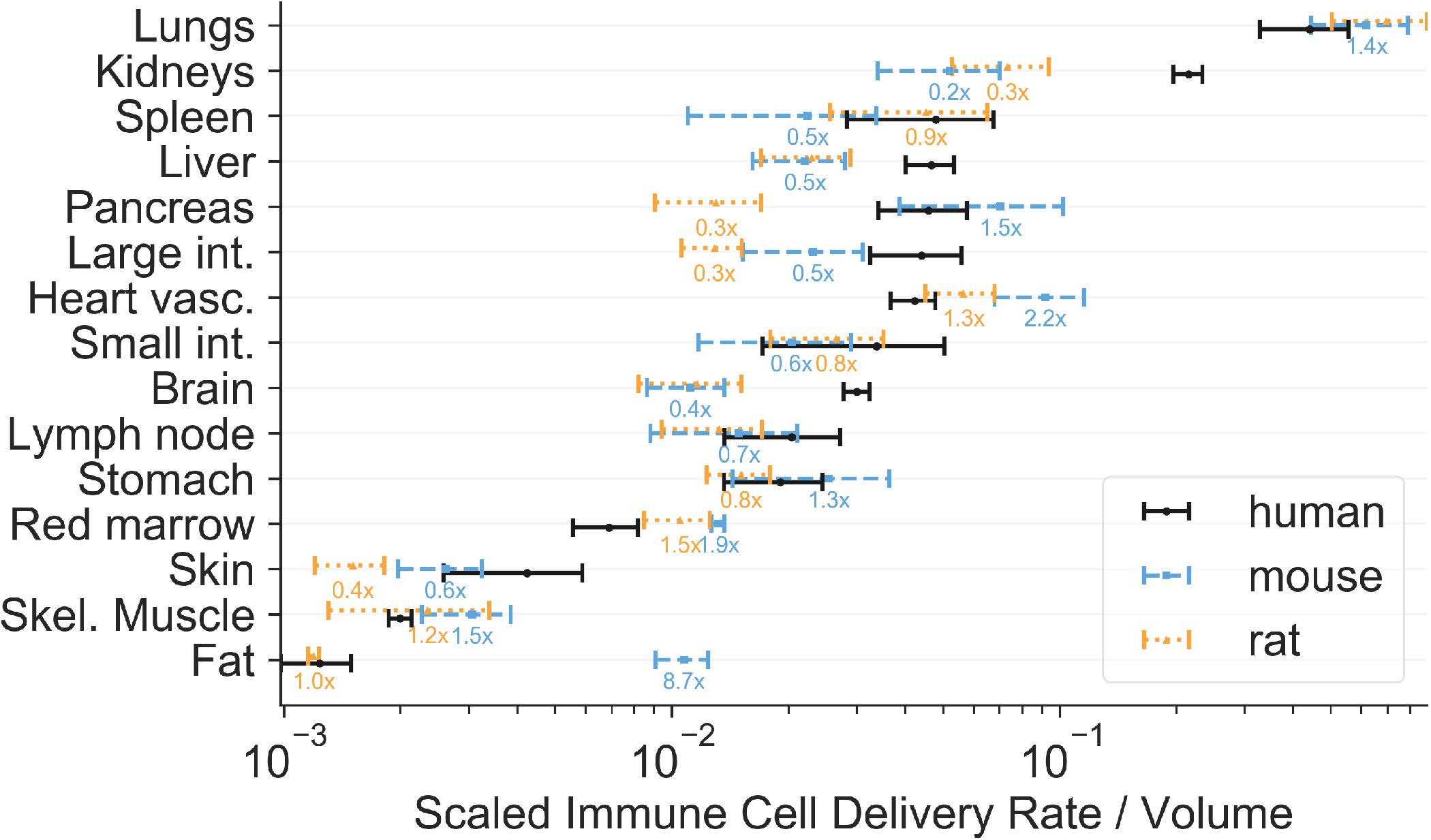
Relative predicted delivery rates to non-tumour tissue in organs in humans, rats and mice. Rates within each species were normalised to sum to 1.0 to give relative values for comparison. See Table 1 for a comparison of absolute rates. Predictions are presented as a mean and standard deviation over 100 repeats, with random fractional blood flows and volumes uniformly drawn from experimental data in the literature. Numerical values (text labels) give the mean predicted delivery rates in rats and mice relative to human values, with colours matching the legend. The relative distribution of rates across the major organs differs by species, and, consequently, inter-species scaling of delivery rates is organ-specific. Note that the horizontal axis is a log scale.

Random results in Figure 3 are obtained similarly, by drawing uniformly random values of organ parameters and normally distributed values of tumour perfusion *P*_tmr_. The maximum delivery to tumour tissue is calculated from 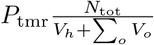 and the maximum delivery rate to non-tumour tissue is calculated using Equation 1 for comparison. As before, *n* = 100 values were chosen for each organ.

**Figure 3:**
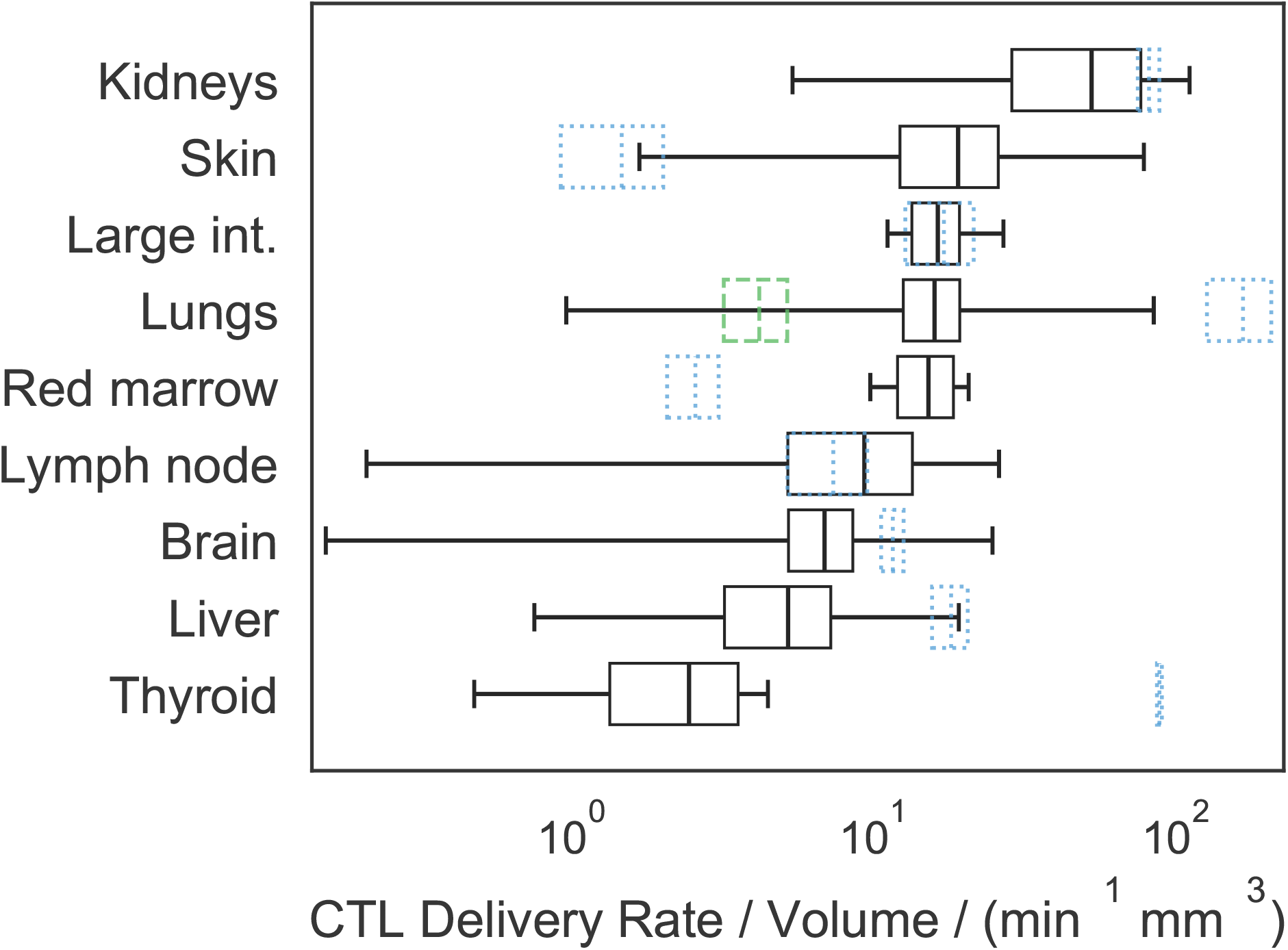
**Absolute predicted delivery rates to human tumours, compared to non-tumour tissue**, assuming 10^8^ CART-cells are administered IV. Predictions are presented as a mean and standard deviation over 100 repeats, with random anatomical parameters and tumour perfusion drawn from experimental data in the literature. Black boxes represent the mean and standard deviation of predicted tumour delivery rates, and whiskers indicate predictions using the extremes of possible tumour perfusion according to the literature. Blue dotted boxes indicate the mean and standard deviation of predicted delivery to non-tumour tissue, *i.e.* the data used to generate Figure 2. The green dashed box indicates delivery rates per mm^3^ to healthy lung tissue when the pulmonary circuit is assumed not to contribute. Note that now the kidneys and skin have the highest predicted tumour delivery rates, and that the horizontal axis is a log scale.

## 3 Results

### 3.1 CART-cell delivery to organs in humans, mice and rats

We calculated and compared predictions for the delivery rate per volume (cells/min/mm^3^) of a typical number of CART-cells used in the clinic (10^8^ [27, 28]) to non-tumour tissues in different human, rat and mouse organs. These rates are equal to the product of the organ perfusion and CART-cell blood concentration. Results calculated from a single anatomical data set ([24]) are shown in Table 1. Flow from both the hepatic artery and portal vein are included in delivery rates to the liver, and the pulmonary circuit and lung blood supply are both included for lung rates. The difference in delivery rates to the same organ in different species can be extreme, with predicted absolute lung delivery rates per volume in the mouse 21,000 times higher than in humans if the same number of CART-cells is administered to each species (obtained by dividing 3,700,000/180 from Table 1). Should a known blood concentration of endogenous cells be considered instead of a constant number, then rates per volume depend only on organ perfusion, and the absolute delivery rates for mice are up to 10 times higher than in humans. These data suggest that a more appropriate approach for scaling murine dosages to humans is to ensure that the same cellular delivery rate to tissues of interest is achieved. The results of Table 1 were used to calculate the CART-cell doses (introduced cell numbers) required to obtain the same delivery rates in humans as in mice given a typical pre-clinical dose of 10^7^ CART-cells. Equivalent doses are organ-specific, and most are of order 10^10^ to 10^11^ cells (Table 2).

The mean and standard deviation of predicted delivery rates obtained by random selection of anatomical parameters from all data sets [21–26] are plotted in Figure 2. To illustrate organ-specific scaling and to allow interspecies comparison of the distribution of delivery rates across organs, rates are normalised such that the sum of the mean predictions within each species is 1.0. The distributions share similarities but otherwise the relative rates exhibit organ-specific scaling. For each species, the lung has the highest delivery rate, followed by the kidneys.

### 3.2 CART-cell delivery to human tumours

Predicted maximum delivery rates per mm^3^ of tissue described above assume that perfusion is homogeneous within a given organ. However, a tumour may have perfusion different to normal tissues. The literature was surveyed to quantify the variability of human tumour perfusion (supplementary figure S3) for incorporation into estimates of maximum delivery rates. As before, delivery rates were calculated with many random values of parameters (*n* = 100 per organ), drawn uniformly for all organ parameters and from a Gaussian distribution for tumour perfusion. The mean and standard deviation of predicted delivery rates for CART-cells to human tumours are shown in Figure 3, along with the corresponding delivery rates under the assumption of homogeneous perfusion (or equivalently, to non-tumour tissue; blue dotted boxes). The rank order of delivery rates to tumour and normal tissues are very different. In most cases, the average of predicted delivery rates for tumour tissue is similar to or less than that for normal tissue, but in some cases (*e.g.* the skin) it is considerably greater. However, their variation is considerable; extreme values (whiskers in the plot) vary over many orders of magnitude above and below that of the corresponding normal tissue, for most organs.

### 3.3 Maximum delivery estimates are consistent with PET imaging and radiography data

The validity of “maximum delivery rates” to organs can be tested by comparing data from PET imaging and radiography studies in humans and rodents, in which cell localisation at early time points has been recorded. The use of an early time point is critical, as it shows the location of cells that are still in the blood or recently extravasated into an organ, before they drain back into the blood and recirculate. At later time points, localisation is a function of both cell delivery to organs and return to circulation. The delivery of radiolabelled natural killer cells from the bloodstream into individual organs has been studied in rats [29] and in human patients [30, 31]. These data are presented in Figure 4 and compared to predictions from Table 1. Patients in the human study were given 10^8^ to 10^9^ cells; the average fraction found in the liver at the first time point (30 minutes) was 8.9%. This corresponds to approximately 4.5×10^7^ cells. The rats were given 10^6^ to 10^7^ cells; the average fraction found in the liver at the first time point (30 minutes) was 23.0%, or 1.2×10^6^ cells. Adjusting the rat numbers to the human dose gives 1.2×10^8^ cells. If we then assume a liver volume of 1700ml in humans and 10ml in rats, we obtain cell number per unit volume in the liver: 2.6×10^4^ in humans and 1.1×10^7^ in rats, a ratio of 429. The ratio of predicted maximum delivery rates is 546 (Table 1), 1.27-fold larger than expected from the data. Repeating this analysis for the lungs and spleen gives experimental ratios 2.0-fold less than predicted from maximum delivery rates (see Figure 4).

**Figure 4:**
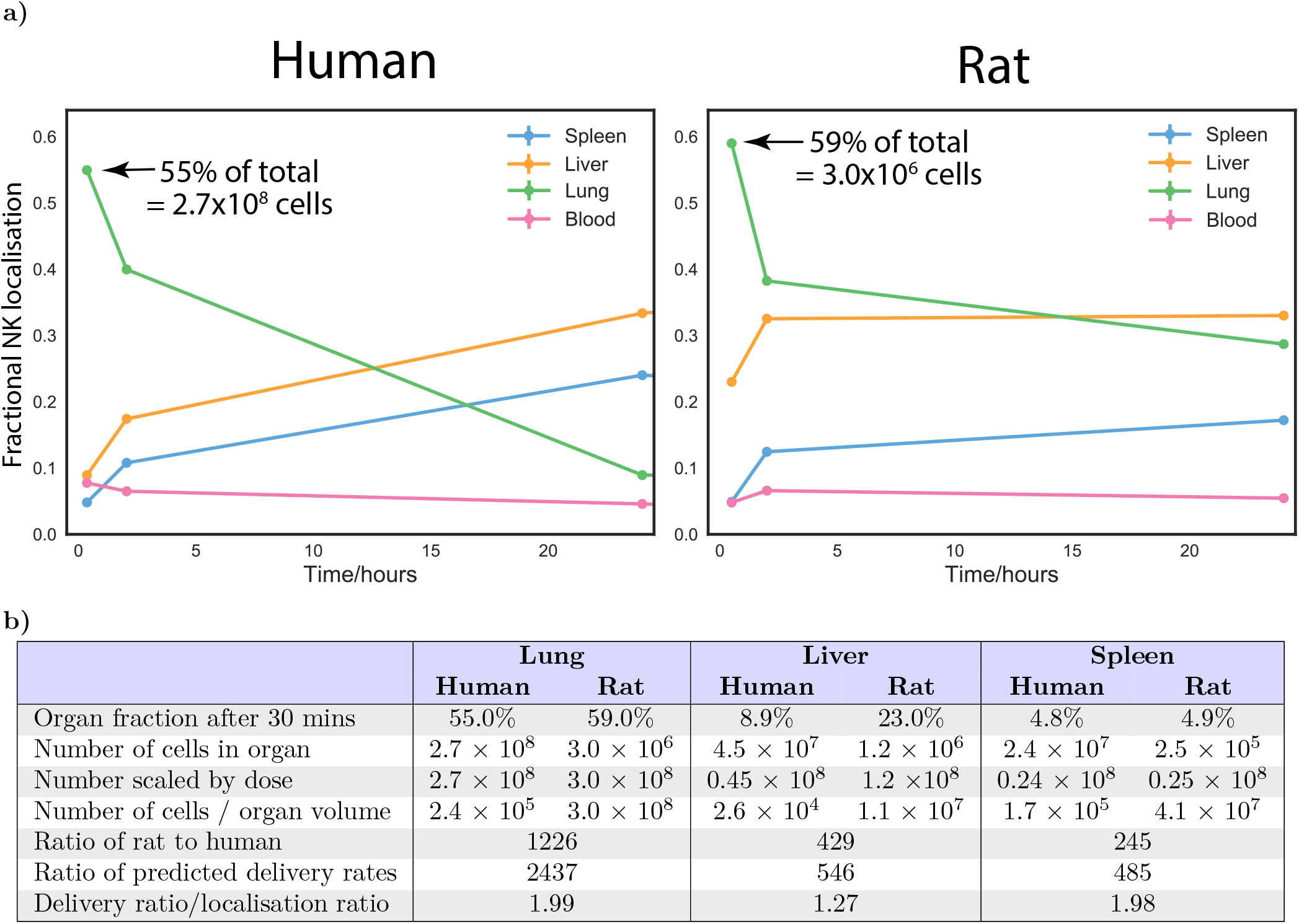
**Comparison of reported localisation of radiolabelled natural killer (NK) cells** in rats and humans to predicted maximum delivery rates [29–31]. **a)** Reproductions of the reported data, after normalising data at each time point such that the total radioactivity (localisation) is 1.0 at all time points. Annotations indicate the initial count of cells in the lung in each species. **b)** Analysis of the data. The dosage and fractional localisation in each organ can be used to calculate the number of NK cells present in each organ at each time point. By accounting for the different dose given to each species and choosing an appropriate estimate for organ volumes in each species, the number of cells per volume in each species can be calculated. The rat/human ratio of the number of cells in each organ can be compared to the ratio of predicted maximum delivery rates per volume, obtained from Table 1.

## 4 Discussion

### 4.1 Vascular delivery and cell proliferation

This study aimed to quantify physiological constraints on the rate of CART-cell delivery by the blood to target tissues in different species, to better predict appropriate clinical CART-cell doses from pre-clinical data. It has focused on adoptive T-cell cancer therapies, though the methodology may also apply to other therapeutic areas, including immune-related adverse event prediction. Values were calculated assuming that 10^8^ T-cells are introduced; delivery rates due to any other desired number or blood concentration of cells can be calculated by multiplying results by the ratio of the desired number to 10^8^ or multiplying blood concentration by the total blood volume in the target species. Although models to predict expansion of a T-cell population have been studied in the past [9, 32], it is difficult to quantify cellular proliferation in or fractional recirculation from a given tissue. However, proliferation itself depends on exposure of transferred T-cells to their target antigen, so early responses are expected to be constrained by delivery. Several studies have established a relationship between dose and response for cellular therapies, despite proliferation increasing the effective dose over time [28, 33–35]. Furthermore, delivery of cells that proliferate outside of a given tumour site would also be constrained by vascular delivery. The maximum rate of delivery due to the anatomy can be estimated with greater confidence and wider applicability than can an estimated time-course of T-cell concentration that considers proliferation and contraction, so proliferation was not considered in this work and will be the focus of future studies.

### 4.2 Organ-specific delivery rates and their variation

Results predict that the highest CART-cell delivery rates are in organs with the highest perfusion: the lungs and kidneys in humans (Figure 2). When measurements of tumour-specific perfusion are considered (Figure 3), it is the kidneys, skin, large intestine and lungs that are predicted to have the highest delivery rates per mm^3^, consistent with non-cellular immunotherapies (IL-2 and checkpoint blockade) having the highest efficacy in kidney, skin, colon and lung tumours [36–41], and the hypothesis that efficacy is driven in part by tissue perfusion. For cellular therapies including CART-cells, vascular delivery should similarly correlate with efficacy, with the additional factor that T-cells must extravasate into target tissues. Both naïve and *ex vivo* T-cells preferentially extravasate into lymph nodes, spleen and liver [42–44], consistent with CART-cell efficacy in haematological disorders but not solid tumours [1, 27, 28]. Tumour tissue may be vascular or avascular, and inflamed or uninflamed, meaning that predicted ‘maximum’ delivery rates for normal tissues may not be applicable to tumours. However, predicted mean delivery rates into tumours exceed those to normal tissue for only a minority of organs (Figure 3), including the skin. Predicted delivery rates to tumours in the skin vary over many orders of magnitude but are usually greater than those for normal tissue. Healthy skin is not usually highly perfused and contains shunts to control blood flow in response to temperature. Most anatomical data for the skin describes the organ at rest and at room temperature with no inflammation, meaning most shunts will be open. Tumour tissue can increase its perfusion through inflammation or angiogenesis and likely subverts these shunts, which could explain the greater mean and variation in predicted delivery rates for skin tumours. Liver and kidney tissues are highly perfused at rest, which are unlikely to be improved by random tumour angiogenesis; accordingly, predicted delivery rates to tumours in these organs do not exceed normal tissue. Predictions for red bone marrow indicate that tumour perfusion can greatly outstrip normal tissue perfusion. Though surprising, the bone red bone marrow result is consistent with studies in which bone perfusion was measured in healthy control bone and tumour sites in patients with bone cancers and metastases [45]. Finally, predicted delivery rates to lung tumours may or may not exceed that of normal tissue, depending on whether the pulmonary circuit is assumed to contribute to tumour blood supply (blue dotted box) or not (green dashed box). Aside from these exceptions, results suggest that predicted maximum delivery rates to normal tissue are greater than those to tumour tissue of the same origin in most cases, and so appropriate to use as a guideline to compare species.

Both figures 2 and 3 show that predicted delivery rates are highly variable, which may be caused by differences in experimental techniques or individual variation. Physiological differences and behaviour both impact blood flow distributions; blood flow to the mesentery increases after a meal, muscles during exercise, or the skin in response to temperature. This effect is utilised in the clinic to prevent hair loss in chemotherapy patients by cooling the scalp. CART-cell therapies could be targeted to organs such as the mesentery or skin through meal consumption or temperature control, and tumour-specific blood flow could be increased with vessel normalisation associated with anti-angiogenic therapies (*e.g.* Avastin). Both human and rodent anatomical parameters vary, impacting any results that depend on anatomical parameters. If variability is not captured and/or care is not taken to control factors that alter blood flows (*e.g.* anaesthesia, exercise or the time of day [46]), then comparison of data sets may be invalid. Ideally, any study making use of blood flows and organ volumes should consider multiple measurements and include ‘error’ bars to indicate variation.

### 4.3 Species-specific delivery rates and dosage scaling

Relative delivery rates are distributed differently across organs in each species, meaning that dose scaling is organ-specific (Figure 2, Table 1). Predicted absolute delivery rates of the same dose of CART-cells (10^8^) exhibited surprisingly extreme differences between species, with delivery per unit tissue volume to mouse lungs 21,000 times higher than in humans, largely because of the difference in total blood volumes between mice (2mL) and humans (5L). To test the relevance of these “maximum delivery rates” and validate the model, we analysed published PET imaging and radiography studies of natural killer (NK) cells in humans and rats [29–31] and calculated the cell numbers present in various organs at early time points (Section 3.3). The human/rat ratios of NK unit volume in the lungs, liver and spleen 30 minutes after infusion were compared to the human/rat ratios of predicted maximum delivery rates. The measured localisation ratios are 1.3 to 2.0-fold greater than predictions for delivery rate ratios. Such small discrepancies are not unexpected, as delivery *rate* ratios would only equal localisation ratios if the blood concentration of NK cells and hence delivery rates were constant. However, the earliest experimental time point is 30 minutes, providing sufficient time for blood recirculation (as cardiac output/minute is greater than total blood volume in humans and rats). The rates of extravasation and return in each organ may differ between humans and rats, and the experimental technique and total amount of radioactivity at the first time point differs between the two studies. Regardless of these potentially confounding factors, the observations are consistent with predictions.

Despite the considerably greater delivery rates of cells in mice than humans, typical doses (cell numbers) introduced to mice are not considerably lower than those given to humans. Most patients are given CART-cell dosages between 10^7^ and 10^9^ cells [27, 28], whilst mouse studies have used (for example) two doses of 1 to 2.5×10^6^ cells a week apart [6], two doses of 10^7^ cells a week apart [3], and a single dose of 10^7^ cells [4]. To illustrate how large these doses are, we calculated equivalent human dosages that would yield the same absolute delivery rates in humans as in a mouse given 10^7^ CART-cells (Table 2). The resulting doses range between 10^10^ and 10^11^ T-cells, much higher than typical clinical doses and many dose escalation studies [28]. This may explain why pre-clinical success does not always translate to the clinic. A pre-clinical study of a CEA CART-cell therapy resulted in regression of subcutaneous tumours in mice with a dose of 5×10^6^ cells (equivalent to 1.7×10^10^ in humans) [47]. In another study, a CART-cell therapy restricted the growth of pancreatic tumours in all treated mice to below the limit of detection with a dose of 10^7^ cells (equivalent to 4.6×10^10^ in humans) [48]. A study in which lower doses of around 10^5^ anti-CD19 cells (human equivalent, using total blood volume only, of 2×10^8^ cells) were given to mice as a ‘stress-test’ was associated with poor tumour control [35]. In the clinic, a study of CEA CART-cells against colorectal cancer [34] escalated doses between 10^7^ and 10^10^ cells. The authors found that the lower doses did not stop tumour progression (in 3 of 14 of presented patients) and higher doses achieved only stable disease. Our results suggest that dosages of order 10^10^ cells would be required to drive tumour regression at the primary site, and 10^11^ would be required for the lung metastases. Clinical studies in which Tumour Infiltrating Lymphocytes (TILs) were introduced in greater numbers (10^9^ to 10^11^) [49–52] and in which CART-cells were introduced regionally (bypassing trafficking via the bloodstream) [27] are associated with greater efficacy. An important caveat of these simple comparisons is that some of the studies lymphodepleted the mice or patients before infusing T-cells, which aids proliferation, and some did not.

In patients with advanced metastatic disease, CART-cell dosage must be sufficient to drive tumour regression at the least perfused and/or the fastest growing site. To avoid dosage-linked increases in adverse events such as cytokine release or encephalopathy syndromes, methods to increase the effective dose on-site and not elsewhere should be considered, including alternate modes of administration, triggering proliferation at sites of interest, coadministration of inhibitors (e.g. anti-IL6), or interventions to alter blood flows described in Section 4.2.

We used natural killer cell localisation data to validate the model, by confirming that early localisation of cells correlates with predicted maximum delivery rates and assuming that natural killer and T-cells behave similarly to each other at short time scales. A more appropriate validation would compare predictions to the localisation of adoptively transferred cells to solid tumours in mice and humans, however, such data is sparsely published and we have found no reported data for humans that includes organ and tumour localisation at an early time point (of the order of minutes). Such data would be useful for further work, as would a time course that could be used to quantify the subsequent constraints imposed by homing and proliferation of cells.

The numbers presented here compare organs like-for-like between mice and humans, but scaling is more uncertain for xenografts. The ratio of the maximum delivery rate per volume to skin tissue between mice and humans is 2 if the same blood concentration of immune cells is assumed, or 3400 if the same number of immune cells is assumed (calculated from Table 1). The ratio of delivery rates per volume to mouse skin versus human kidney tissue, meanwhile, is 0.05 if the same concentration of cells is used, or 100 if the same number of immune cells is assumed. A previous study [53] has shown that small xenografts have similar local perfusion to the original tissue, but larger xenografts have reduced perfusion relative to the original tissue. This non-linearity further confounds extrapolation of preclinical results. This highlights some of the historically observed difficulties in the clinical translation of preclinical mouse xenograft model results [54]. In addition to consideration of physiological and immunological differences (such as the adhesion molecules required for T-cell extravasation), interpretation of pre-clinical therapeutic success requires dosages to be appropriately scaled to humans. A model that considers organ-specific blood flow and volumes across species can be used to more precisely estimate likely efficacious human doses.

### 4.4 Prediction refinement by T-cell homing and further considerations

The presented results are the predicted maximum delivery rates of CART-cells per unit volume (cells/min/mm^3^) to organs and tumours, based on only organ blood flows and volumes. Refining these predictions requires quantification of CART-cell proliferation and organ-specific homing. The probability of T-cell extravasation differs by location and cell type. Naïve T-cells extravasate mainly into the lymph nodes or spleen and activated cells have a higher probability of extravasating into non-lymphoid tissues [42, 55], distributed according to upregulated homing receptors (*e.g.* L-selectin or CCR7 [56]). These probabilities may differ across species (*e.g.* homing receptor CXCR1 is present in humans but not mice [57]), further limiting inter-species extrapolation of pre-clinical results. Homing receptor density, vessel normalisation and hence homing probabilities may further differ in tumour tissue, particularly following therapies such as Avastin [58, 59]. It is possible to quantify organ-specific homing by fitting ODE models (like the model shown in supplementary section A.1) to T-cell localisation data in experimental animals, as previous authors have done, *e.g.* [19]. However, we have found very limited equivalent human data for cross-species comparison, which is the primary aim of this work. Parameters obtained from fits to multiple experiments would differ due to differences in the animals and the cells, so several datasets would be required to quantify the variation of and/or a confidence interval for parameter estimates. The focus of this study is on anti-tumour therapies, where tumour homing would be affected by factors such as inflammation. For this reason, we chose to quantify *maximum* delivery rates by examining the case where T-cells have a 100% probability of extravasation in the target organ, and no extravasation elsewhere. Species comparisons are made by implicitly assuming that homing probabilities in a given organ or tumour would be similar between species. Expected variation in predictions was quantified by using the variation among anatomical reference values as a proxy. Both maximum values and this variation could be improved by more precise measurements of blood flows and volumes using the same techniques in each species, or else finding anatomical parameters for a precise experimental animal of interest.

Another challenge for CART-cells in solid tumours is the identification of suitable target antigen. The ideal antigen is highly expressed on tumour cells and not expressed on healthy cells elsewhere. A typical target for B-cell malignancies is CD19 [28], as it is expressed by the entire pool of B-cells and is limited almost exclusively to B-cells. Several different antigens have been targeted for solid tumours, but with limited success (for example, GD2 has had encouraging results [28]). Target antigen may only be expressed by a subset of tumour cells and may not be sufficiently rare elsewhere in the body. For example, CAIX is expressed in some renal cell carcinomas, but it is also expressed in the liver bile duct resulting in on-target, off-tumour toxicities in a phase III trial [60]. Tumours may evolve to reduce expression of target antigen in response to successful T-cell killing, reducing the rate of tumour elimination or promoting outgrowth of therapy-resistant cells. Although these considerations are a barrier to treatment success, the rate at which cells can be delivered is a parallel and important factor. CART-cells that are specific for an antigen that is expressed on most tumour cells will not drive tumour regression if their kill rate is lower than the tumour growth rate, given the combined rates of T-cell delivery and proliferation. On the other hand, CART-cells specific for a rarer antigen may drive tumour regression if they arrive in sufficient numbers to eliminate all cells carrying that antigen, subsequently proliferating to greater numbers to drive regression at more restricted sites and/or drive a secondary response against one or more other antigens (*i.e.* epitope spread). Like T-cell delivery rates and T-cell extravasation probabilities, typical tumour growth rates are species, organ and individual specific. Together, these considerations show that tumour immunotherapy is a numbers game and hence more generally quantitative studies can be a useful tool for understanding the translational gap between pre-clinical and clinical outcomes.

## 5 Conclusions

Details of the human, rat and mouse circulatory systems were considered to predict CART-cell delivery to human tumours, and to human, rat and mouse organs. Predictions show up to an order of 10,000-fold increased CART-cell delivery per unit volume of target tissue in mice than humans, while typical clinical cell therapy dosages are 100-fold less than typical pre-clinical doses. These numbers are consistent with experimental studies of NK cell localisation and various clinical observations. These predictions could partially explain why pre-clinical models of solid tumour clearance by CART-cells show greater efficacy than in humans. Dosage scaling was found to be organ-specific and is particularly hard to quantify for xenografts, confounding the interpretation of pre-clinical results and lowering their potential clinical value, which is an important consideration in the context of the reduction and replacement of animal experiments. Control of tumour and organ-specific blood flow through exercise, circadian timing or food consumption could increase cellular delivery to tumour sites without raising the prospect of adverse outcomes, while vascular normalisation may also induce such benefits, though with accompanying risk. More generally, cellular kinetic and dynamic models will lead to better understanding of how pre-clinical outcomes translate to the clinic, and hence better determination of appropriate clinical dosages and treatment strategies for cell-based therapies.

## Declarations

### Availability of data and code

Code and parameter data for humans, mice and rats used to generate the results of this work are available as supplementary materials.

### Author contributions

LVB designed the model code, did the analysis and wrote the manuscript. All other authors supervised, advised and edited the manuscript.

