## Supplementary Material for "Quantifying the Limits of Immunotherapies in Mice and Men"

March 2, 2020

### A Methods

#### A.1 ODE model definition

As stated in the main text, many studies of physiologically-based pharmacokinetics or cellular kinetics make use of systems of ordinary differential equations (ODEs), which represent the exchange of material between the blood and organs. The results of the main text can be produced from such a system, rather than the simpler treatment using the expression *perfusion* × *concentration* that was presented. The construction of an equivalent model and results with ODEs is presented here to show consistency and parity with existing literature, and to assist other authors with extension of this work using a more complete modelling framework.

##### A.1.1 ODE system equations

| Symbol | Definition |
|---|---|
| $V_o$ | blood volume in organ $o$ |
| $B_o$ | blood flow to organ $o$ |
| $C_o$ | (CAR) T-cell blood concentration (1/mm$^3$) in organ $o$ |
| $\tilde{V}_o$ | total volume of organ $o$ |
| $\tilde{C}_o$ | interstitial cell concentration in organ $o$ |
| $e_o$ | proportion of cells extravasating to interstitial space (such that $e_o B_o C_o$ is the entry rate) |
| $\mu_o$ | proportion of cells leaving the interstitial space for the lymph node ($e_o \mu_o B_o \tilde{C}_o$ is the exit rate) |

**Table S1. Definition of parameters** described in supplementary section A.1. Different suffixes such as $h$ for heart, $\mathrm{tmr}$ for the tumour and $\mathrm{TBO}$ for the tumour-bearing organ are also used.

The circulatory system can be implemented as a system of ODEs representing material exchange between the heart and organs. The change in concentration of immune cells in the vasculature of any organ $C_o$ is given by,

$$V_o \frac{\mathrm{d}C_o}{\mathrm{d}t} = B_o(C_\mathrm{h} - C_o), \tag{1}$$

where $V_o$ and $B_o$ are constant and are respectively the blood volume and the blood flow of the organ, whilst $C_\text{h}$ is the immune cell concentration within the heart chambers. A proportion of cells leaving the organ $e_o B_o$ extravasate to the interstitial space:

$$\tilde{V}_o \frac{\mathrm{d}\tilde{C}_o}{\mathrm{dt}} = e_o B_o \left(C_o - \mu_o \tilde{C}_o\right) \tag{2}$$

where $\tilde{V}_o$ is the total volume of the organ, $\tilde{C}_o$ is the interstitial concentration and $e_o \mu_o B_o$ is the flow of immune cells into the lymph from the interstitium of organ $o$. Note that $\tilde{V}_o$ has not had the vascular volume subtracted from it, to avoid the combination of measurements from different experiments in a single organ's data, and to avoid cases where the interstitial volume is not well-defined or certain. The percentage error from this approximation is small (see supplementary tables S3 and S4). Both equations 1 and 2 are used for all organs, save a few exceptions. For example, both equations are modified for the liver, which also receives blood exiting the mesenteric organs:

$$\begin{aligned}
V_\text{liver} \frac{\mathrm{d}C_\text{liver}}{\mathrm{dt}} &= B_\text{liver}(C_\text{h} - C_\text{liver}) + e_\text{spleen}\mu_\text{spleen} B_\text{spleen}(\tilde{C}_\text{spleen} - C_\text{liver}) + \sum_{o' \in \text{mesentery}} B_{o'}(C_{o'} - C_\text{liver})(1 - e_{o'}) \\
\tilde{V}_\text{liver} \frac{\mathrm{d}\tilde{C}_\text{liver}}{\mathrm{dt}} &= e_\text{liver} B_\text{liver} \left(C_\text{liver} - \mu_\text{liver}\tilde{C}_\text{liver}\right) \\
&\quad + \sum_{o' \in \text{mesentery}} e_\text{liver} \left(C_\text{liver} - \mu_\text{liver}\tilde{C}_\text{liver}\right) B_{o'}(1 - e_{o'}) \\
&\quad + e_\text{liver} \left(C_\text{liver} - \mu_\text{liver}\tilde{C}_\text{liver}\right) e_\text{spleen}\mu_\text{spleen} B_\text{spleen},
\end{aligned} \tag{3}$$

where the summed terms arise from the extravasation and return of cells that did not extravasate in the mesentery and instead flowed through the portal vein, and the final line represents cells returning from the spleen. Cells entering the lymphatics flow towards the lymph nodes:

$$\begin{aligned}
V_\text{LN} \frac{\mathrm{d}\tilde{C}_\text{LN}}{\mathrm{dt}} =& e_\text{LN} B_\text{LN} \left(C_\text{LN} - \mu_\text{LN}\tilde{C}_\text{LN}\right) \\
&+ \sum_{o' \notin \{\text{LN,spleen,PC}\}} e_{o'}\mu_{o'} B_{o'}(\tilde{C}_{o'} - \mu_\text{LN}\tilde{C}_\text{LN}) \\
&+ e_\text{liver}\mu_\text{liver} \left( e_\text{spleen}\mu_\text{spleen} B_\text{spleen} + \sum_{o' \in \text{mesentery}} B_{o'}(1 - e_{o'}) \right) (\tilde{C}_\text{liver} - \mu_\text{LN}\tilde{C}_\text{LN}),
\end{aligned} \tag{4}$$

where the second line represents the flow of lymphocytes from other organs, the third line is additional flow from the liver, and PC is the pulmonary circuit. Cells that do not extravasate from the organ vasculature return to the heart, in addition to the lymph node output:

$$\begin{aligned}
V_\text{h} \frac{\mathrm{d}C_\text{h}}{\mathrm{dt}} =& - C_\text{h} \sum_o B_o \\
&+ \sum_{o \notin \text{mesentery}} (1 - e_o) B_o C_o \\
&+ \left( e_\text{spleen}\mu_\text{spleen} B_\text{spleen} + \sum_{o \in \text{mesentery}} (1 - e_o) B_\text{o} \right) (1 - e_\text{liver}) C_\text{liver} \\
&+ e_\text{LN}\mu_\text{LN} B_\text{LN}\tilde{C}_\text{LN} + \sum_{o \notin \{\text{spleen,PC,LN}\}} \mu_\text{LN} e_o \mu_o B_o \tilde{C}_\text{LN} \\
&+ e_\text{liver}\mu_\text{liver}\mu_\text{LN}\tilde{C}_\text{LN} \left( e_\text{spleen}\mu_\text{spleen} B_\text{spleen} + \sum_{o \in \text{mesentery}} B_o \left(1 - e_o\right) \right) \\
&+ e_\text{PC}\mu_\text{PC} B_\text{PC}\tilde{C}_\text{PC},
\end{aligned} \tag{5}$$

where each line respectively refers to: (i) movement of cells out of the heart, (ii) return of cells from non-mesenteric organs, (iii) return of cells from the mesentery via the liver, (iv) return of cells from the lymph nodes, (v) extra return of cells from the liver via the LN, and (vi) return of cells from the pulmonary circuit.
Note that $V_h$ is the blood volume of the heart chambers. Venous and arterial compartments are not modelled and

are not counted in the total blood volume, as transit through them is assumed to be rapid compared to transit through organs (venules and arterioles). We may designate an organ as a tumour-bearing organ (or any other disease of interest), and define the rate of immune cell entry to the tumour as,

$$\frac{\mathrm{d}N_{\mathrm{infiltrate}}}{\mathrm{dt}} = e_{\mathrm{tmr}} \frac{\tilde{V}_{\mathrm{tmr}}}{\tilde{V}_{\mathrm{TBO}}} B_{\mathrm{TBO}} C_{\mathrm{TBO}}, \tag{6}$$

where $\tilde{V}_{\mathrm{tmr}}$ is the tumour volume and $\tilde{V}_{\mathrm{TBO}}$ the organ volume of the tumour-bearing organ. Extravasation to the rest of the organ is reduced due to the effective loss of organ volume and hence:

$$\tilde{V}_{\mathrm{TBO}} \frac{\mathrm{d}\tilde{C}_{\mathrm{TBO}}}{\mathrm{dt}} = e_{\mathrm{TBO}} \left( C_{\mathrm{TBO}} - \mu_{\mathrm{TBO}} \tilde{C}_{\mathrm{TBO}} \right) \left( 1 - \frac{\tilde{V}_{\mathrm{tmr}}}{\tilde{V}_{\mathrm{TBO}}} \right) B_{\mathrm{TBO}}. \tag{7}$$

The flow of cells that do not return to the heart from the TBO is subsequently changed from $B_{\mathrm{TBO}} C_{\mathrm{TBO}} e_{\mathrm{TBO}}$ to

$$B_{\mathrm{TBO}} C_{\mathrm{TBO}} \left( e_{\mathrm{TBO}} \left( 1 - \frac{\tilde{V}_{\mathrm{tmr}}}{\tilde{V}_{\mathrm{TBO}}} \right) + e_{\mathrm{tmr}} \frac{\tilde{V}_{\mathrm{tmr}}}{\tilde{V}_{\mathrm{TBO}}} \right),$$

with changes for the liver and lymph nodes made analogously to before. Note that tumour-invading lymphocytes are assumed never to return via the lymphatics, and that these equations implicitly assume that tumour perfusion is equal to that of the rest of the organ, without modification.
The total number of ODEs is $2+2+2n$, where $n$ is the number of organs in the system. The first '2' is the equations for the heart and tumour, the second is an extra pair of compartments for the lung pulmonary circuit with a trapped state for lymphocytes, and $2n$ is the vascular and interstitial compartments for all other organs. There are three fixed parameters per organ (the fractional blood flow and fractional blood volume, and organ volume, where fractional parameters sum to 1), and three system-wide fixed parameters (the cardiac output, the total blood volume and the dosage of T-cells). These values are constrained by our anatomical knowledge or known input, so are not varied. There are two unknown parameters per organ (the extravasation probability $e$ and return rate $\mu$).

##### A.1.2 Including tumour perfusion in the ODE model

If tumour perfusion (blood flow per volume) is expected to be different from the perfusion of the healthy organ, then we define new blood flows $B'_{\mathrm{tmr}}$ and $B'_{\mathrm{healthy}}$ such that $B'_{\mathrm{TBO}} = B'_{\mathrm{tmr}} + B'_{\mathrm{healthy}}$ and their perfusion ratio is,

$$p := \frac{B'_{\mathrm{tmr}} / \tilde{V}_{\mathrm{tmr}}}{(B'_{\mathrm{TBO}} - B'_{\mathrm{tmr}}) / (\tilde{V}_{\mathrm{TBO}} - \tilde{V}_{\mathrm{tmr}})}. \tag{8}$$

Note that $\tilde{V}_{\mathrm{tmr}} \lesssim \tilde{V}_{\mathrm{TBO}}$ and $B_{\mathrm{tmr}} \lesssim B_{\mathrm{TBO}}$ (that is, the volume and blood flow of the tumour bearing organ explicitly include those of the tumour), so the perfusion ratio is strictly positive. Equation 8 may be used to calculate an expression for the new tumour blood flow in terms of $p$,

$$B'_{\mathrm{tmr}} = B'_{\mathrm{TBO}} \frac{p \frac{\tilde{V}_{\mathrm{tmr}}}{\tilde{V}_{\mathrm{TBO}}}}{1 + \frac{\tilde{V}_{\mathrm{tmr}}}{\tilde{V}_{\mathrm{TBO}}} (p-1)}. \tag{9}$$

To modify tumour perfusion in this way whilst keeping the total blood flow constant, all other blood flows must be reduced. This reduction can be derived as follows, where $\Delta P$ and $\Delta P'$ are the pressure drop across the heart with and without the tumour, $R_o = \Delta P / B_o$ is the vascular resistance to blood flow in organ $o$, the healthy part of the tumour bearing organ is counted within the sum of organ blood flows $\sum_o B_o$, and the total cardiac output $B_{\mathrm{tot}}$

and all vascular resistances other than that of the TBO are assumed to be constant:

$$
\begin{aligned}
B_{\text{tot}} &= \sum_o B_o = \sum_o B'_o = \Delta P' \left( \sum_{o \neq \text{tmr}} \frac{1}{R_o} \right) + B'_{\text{tmr}} \\
\therefore \Delta P' &= \left( B_{\text{tot}} - B'_{\text{tmr}} \right) / \left( \sum_{o \neq \text{tmr}} \frac{1}{R_o} \right) \\
B'_o &= \frac{\Delta P'}{R_o} = B_o \frac{\Delta P'}{\Delta P} \\
&= \frac{B_o}{\Delta P} \left( B_{\text{tot}} - B'_{\text{tmr}} \right) / \left( \sum_{o \neq \text{tmr}} \frac{B_o}{\Delta P} \right) \\
&= B_o \left( B_{\text{tot}} - B'_{\text{tmr}} \right) / \left( B_{\text{tot}} - B_{\text{tmr}} \right) \\
&=: B_o \theta.
\end{aligned} \tag{10}
$$

This allows the blood flow to the healthy part of the tumour bearing organ to be written as

$$
B'_{\text{healthy}} = \theta B_{\text{healthy}} = \theta B_{\text{TBO}} \left( 1 - \frac{\tilde{V}_{\text{tmr}}}{\tilde{V}_{\text{TBO}}} \right) \tag{11}
$$

and the overall blood flow to the tumour bearing organ as,

$$
B'_{\text{TBO}} = B'_{\text{healthy}} + B'_{\text{tmr}} = \frac{B_{\text{tot}} - B'_{\text{tmr}}}{B_{\text{tot}} - B_{\text{tmr}}} B_{\text{TBO}} \left( 1 - \frac{\tilde{V}_{\text{tmr}}}{\tilde{V}_{\text{TBO}}} \right) + B'_{\text{tmr}}. \tag{12}
$$

After substitution of equation 9 and $B_{\text{tmr}} = \frac{\tilde{V}_{\text{tmr}}}{\tilde{V}_{\text{TBO}}} B_{\text{TBO}}$, this can be rearranged to give,

$$
B'_{\text{TBO}} = B_{\text{TBO}} \frac{1 + \frac{\tilde{V}_{\text{tmr}}}{\tilde{V}_{\text{TBO}}}(p-1)}{1 + \frac{\tilde{V}_{\text{tmr}}}{\tilde{V}_{\text{TBO}}}(p-1)\frac{B_{\text{TBO}}}{B_{\text{tot}}}}. \tag{13}
$$

A similar expression can then be derived for $B'_{\text{tmr}}$,

$$
B'_{\text{tmr}} = B_{\text{TBO}} \frac{p\frac{\tilde{V}_{\text{tmr}}}{\tilde{V}_{\text{TBO}}}}{1 + \frac{\tilde{V}_{\text{tmr}}}{\tilde{V}_{\text{TBO}}}(p-1)\frac{B_{\text{TBO}}}{B_{\text{tot}}}} = B_{\text{tot}} \frac{p\frac{\tilde{V}_{\text{tmr}}}{\tilde{V}_{\text{TBO}}}}{\frac{B_{\text{tot}}}{B_{\text{TBO}}} + \frac{\tilde{V}_{\text{tmr}}}{\tilde{V}_{\text{TBO}}}(p-1)}, \tag{14}
$$

which in turn allows $\theta$ to be written in terms of $p$:

$$
\begin{aligned}
\theta &= \frac{B_{\text{tot}} \left( 1 - \frac{p\frac{\tilde{V}_{\text{tmr}}}{\tilde{V}_{\text{TBO}}}}{\frac{B_{\text{tot}}}{B_{\text{TBO}}} + \frac{\tilde{V}_{\text{tmr}}}{\tilde{V}_{\text{TBO}}}(p-1)} \right)}{B_{\text{tot}} - \frac{\tilde{V}_{\text{tmr}}}{\tilde{V}_{\text{TBO}}} B_{\text{TBO}}} \\
&= \frac{B_{\text{tot}}}{B_{\text{tot}} + \frac{\tilde{V}_{\text{tmr}}}{\tilde{V}_{\text{TBO}}}(p-1)B_{\text{TBO}}},
\end{aligned} \tag{15}
$$

which finally allows the blood flows to the tumour and all organs to be redefined in terms of $p$. Note that the total blood flow to the tumour bearing organ, equation 13, is greater ($p > 1$) or less ($p < 1$) than the original blood flow $B_{\text{TBO}}$, implying that growth of a vascular tumour increases overall blood flow to the organ and that the growth of an avascular tumour decreases overall blood flow to the organ.

##### A.1.3 ODE initial conditions

Initial conditions can take one of two forms. The first is a distribution of cells by blood volume,

$$\begin{aligned} V_h C_h(0) =& N_{tot} \frac{V_h}{V_{tot}}, \\ V_o C_o(0) =& N_{tot} \frac{V_o}{V_{tot}}, \\ \tilde{V}_o \tilde{C}_o(0) =& 0, \\ N_{infiltrate}(0) =& 0, \end{aligned} \tag{16}$$

where $N_{tot}$ is the total number of lymphocytes in the system and $V_{tot} = V_h + \sum_o V_o$. The other form is a distribution of cells by blood flow,

$$\begin{aligned} V_h C_h(0) =& N_{tot} \frac{V_h}{V_{tot}} \\ V_o C_o(0) =& N_{tot} \left(1 - \frac{V_h}{V_{tot}}\right) \frac{B_o}{B_{tot}} \\ \tilde{V}_o \tilde{C}_o(0) =& 0 \\ N_{infiltrate}(0) =& 0, \end{aligned} \tag{17}$$

where $B_{tot} = \sum_o B_o$. The relaxation time of the vascular ODE system is fast, so results are not affected by the choice of initial conditions. Were it to matter, the latter choice of initial conditions would be used for injected T-cells (*e.g.* radiolabelled or CART-cells introduced to the bloodstream at time 0), and the former for endogenous immune cells (with uniform blood concentration).

##### A.1.4 Deriving maximum delivery rates from the ODE model

The main results of the text are the maximum delivery rate of cells per volume to organ tissue, *i.e.* perfusion times cell concentration, $\frac{B_o}{\tilde{V}_o} \frac{N_{tot}}{V_h + \sum_o V_o}$ for each organ $o$, where $N_{tot}$ is the total number of lymphocytes in the system, and with some modification due to details of the vasculature; for example, the portal vein blood flow must be added to $B_o$ for the liver (see the analogous modifications to ODEs above). The maximum delivery rate to a tumour with perfusion $P_{tmr}$ distinct from the rest of the organ is obtained by replacing $\frac{B_o}{\tilde{V}_o}$ with $P_{tmr} = \frac{pB_{tot}/\tilde{V}_{TBO}}{\frac{B_{tot}}{B_{TBO}} + \frac{\tilde{V}_{tmr}}{\tilde{V}_{TBO}}(p-1)}$.

Maximum delivery rates per volume may be acquired from the ODE solution by setting $e_{tmr}$ to 1.0 in a 1mm$^3$ volume within an organ of interest and $e_{o'}$ for all other organs $o'$ to 0.0, then recording the rate of entry of cells to the volume of interest at an early time point, after vascular compartments have reached equal concentration. The rate calculated for organ $o$ is insensitive to the values $e_{o'}$ in any other organ $o'$ if the blood concentration of T-cells is constant (see sensitivity analysis in supplementary section B.6). An appropriate early time for vascular dynamics to have steadied with typical blood flows and volumes is at *e.g.* time $t = 10$ minutes with timestep $\Delta t = 1$ minute.

### B Results

#### B.1 Anatomical parameters

Anatomical parameters (fractional blood flow, cardiac output, fractional blood volume, total blood volume and organ volume) sourced from the literature for each species are shown in supplementary tables S2, S3 and S4, along with references.

#### B.2 Predicted absolute delivery rates per volume

Supplementary figure S1 shows the absolute values for predictions in humans. In order to compare the relative rates of delivery between species, these absolute rates were scaled such that the sum in each species was 1.0.

| | H (ICRP) | M (Brown) | H (Peters) | R (Peters) | H (Shah) | M (Shah) | R (Shah) |
|---|---|---|---|---|---|---|---|
| stomach | 0.01 [1] | 0.0237 [2] | 0.00352 [3] | 0.0173 [3] | 0.005 [1] | 0.00965 [3] | 0.0173 [3] |
| smallIntestine | 0.1 [1] | 0.0391 [2] | 0.0819 [3] | 0.110 [3] | 0.0340 [4] | 0.0779 [4] | 0.0676 [4] |
| largeIntestine | 0.04 [1] | 0.0352 [2] | | | 0.0354 [4] | 0.0232 [4] | 0.0268 [4] |
| pancreas | 0.01 [1] | 0.0320 [2] | 0.0120 [3] | 0.00844 [3] | 0.00840 [4] | 0.00836 [4] | 0.0108 [4] |
| spleen | 0.03 [1] | 0.01 [2] | 0.00692 [3] | 0.0106 [3] | 0.0174 [4] | 0.0110 [4] | 0.0304 [4] |
| liver | 0.065 [1] | 0.02 [2] | 0.132 [3] | 0.169 [3] | 0.0363 [4] | 0.0138 [4] | 0.00358 [4] |
| brain | 0.12 [1] | 0.033 [2] | 0.0630 [3] | 0.0112 [3] | 0.0590 [4] | 0.0158 [4] | 0.0111 [4] |
| heart | 0.04 [1] | 0.066 [2] | 0.0135 [3] | 0.0331 [3] | 0.0213 [4] | 0.0489 [4] | 0.0256 [4] |
| lungs | 0.025 [1] | | 0.447 [3] | 0.429 [3] | 0.500 [4] | 0.500 [4] | 0.500 [4] |
| kidneys | 0.19 [1] | 0.091 [2] | 0.0989 [3] | 0.0779 [3] | 0.100 [4] | 0.0918 [4] | 0.0620 [4] |
| fat | 0.05 [1] | 0.09 [2] | 0.0233 [3] | 0.00338 [3] | 0.0309 [4] | 0.0180 [4] | 0.0380 [4] |
| skin | 0.05 [1] | 0.058 [2] | 0.0271 [3] | 0.0422 [3] | 0.0320 [4] | 0.0373 [4] | 0.0340 [4] |
| skeletalMuscle | 0.17 [1] | 0.159 [2] | 0.0674 [3] | 0.0633 [3] | 0.0920 [4] | 0.115 [4] | 0.157 [4] |
| skeleton | 0.05 [1] | 0.11 [2] | 0.0227 [3] | 0.0214 [3] | 0.00712 [4] | 0.0204 [4] | 0.0104 [4] |
| redMarrow | 0.03 [5] | 0.029 [2] | | | 0.03 [5] | 0.029 [2] | 0.021 [2] |
| thyroid | 0.015 [1] | | | | | | |
| gonads | 0.0005 [1] | | 0.000252 [3] | 0.00380 [3] | | | |
| adrenals | 0.003 [1] | | | | | | |
| bladder | 0.0006 [1] | | | | | | |
| lymphNodes | 0.017 [1] | 0.0134 [6–8] | 0.017 | 0.0134 | 0.0101 [4] | 0.00221 [4] | 0.00285 [4] |
| thymus | | | | | 0.000970 [4] | 0.00159 [4] | 0.00207 [4] |
| misc | 0.0139 [1] | 0.220 [2, 6–8] | | | 0.0152 [4] | 0.0146 [4] | 0.0178 [4] |
| **cardiacOutput** | 6.5 [9] | 0.004 [2] | 5.89 [3] | 0.0628 [3] | 5.51 [4] | 0.0113 [4] | 0.0892 [4] |

**Table S2.** Fractional blood flow (H: Human, M: Mouse, R: Rat). Cardiac output is given in litres/minute.

| | H (ICRP) | M (Brown) | H (Peters) | R (Peters) | H (Shah) | M (Shah) | R (Shah) |
|---|---|---|---|---|---|---|---|
| stomach | 0.01 [1] | 0.00857 [2, 8] | 0.00271 [3] | 0.00529 [3] | 0.01 [1] | 0.00523 [3] | 0.00529 [3] |
| smallIntestine | 0.038 [1] | 0.0191 [2, 8] | 0.0283 [3] | 0.0451 [3] | 0.00359 [4] | 0.0142 [4] | 0.0100 [4] |
| largeIntestine | 0.022 [1] | 0.00600 [2, 8] | | | 0.00510 [4] | 0.00612 [4] | 0.00577 [4] |
| pancreas | 0.006 [1] | 0.0159 [2, 8] | 0.00140 [3] | 0.00584 [3] | 0.00333 [4] | 0.00653 [4] | 0.00689 [4] |
| spleen | 0.014 [1] | 0.0213 [2, 8] | 0.00362 [3] | 0.00303 [3] | 0.0156 [4] | 0.0188 [4] | 0.0422 [4] |
| liver | 0.1 [1] | 0.215 [2, 8] | 0.0301 [3] | 0.0463 [3] | 0.107 [4] | 0.201 [4] | 0.169 [4] |
| brain | 0.012 [1] | 0.0435 [2, 8] | 0.0315 [3] | 0.00973 [3] | 0.0186 [4] | 0.0131 [4] | 0.00631 [4] |
| heart | 0.01 [1] | 0.0418 [2, 8] | 0.00525 [3] | 0.00433 [3] | 0.00767 [4] | 0.00716 [4] | 0.00496 [4] |
| lungs | 0.125 [1] | 0.0975 [2, 8] | 0.0206 [3] | 0.00458 [3] | 0.0321 [4] | 0.0361 [4] | 0.0291 [4] |
| kidneys | 0.02 [1] | 0.163 [2, 8] | 0.00555 [3] | 0.0107 [3] | 0.0106 [4] | 0.0353 [4] | 0.0166 [4] |
| fat | 0.05 [1] | 0.0975 [2, 8] | 0.0481 [3] | 0.00865 [3] | 0.0864 [4] | 0.0266 [4] | 0.0458 [4] |
| skin | 0.03 [1] | 0.0761 [2, 8] | 0.119 [3] | 0.162 [3] | 0.0742 [4] | 0.230 [4] | 0.235 [4] |
| skeletalMuscle | 0.14 [1] | 0.184 [2] | 0.575 [3] | 0.617 [3] | 0.386 [4] | 0.305 [4] | 0.337 [4] |
| skeleton | 0.07 [1] | | 0.0939 [3] | 0.0368 [3] | 0.131 [4] | 0.0760 [4] | 0.0581 [4] |
| redMarrow | 0.04 [5] | 0.0277 [10] | | | 0.04 [5] | 0.0277 [10] | 0.021 [2] |
| thyroid | 0.0006 [1] | | | | | | |
| gonads | 0.0004 [1] | | 0.000739 [3] | 0.0140 [3] | | | |
| adrenals | 0.0006 [1] | | | | | | |
| bladder | 0.0002 [1] | | | | | | |
| lymphNodes | 0.002 [1] | 0.0108 [2, 8] | 0.002 | 0.0108 | 0.002 [1] | 0.0108 [2, 8] | 0.0108 |
| thymus | | | | | 0.000206 [4] | 0.000609 [4] | 0.000665 [4] |
| misc | 0.0192 [1] | 0 [2, 8] | | | 0.119 [4] | 0.0239 [4] | 0.0322 [4] |
| **totalBloodVolume** | 5.3 [9] | 0.00146 [2, 11] | 8.386 [3] | 0.0518 [3] | 3.114 [4] | 0.00149 [4] | 0.0144 [4] |

**Table S3.** Fractional blood volume (H: Human, M: Mouse, R: Rat). Total blood volume is given in litres.

| | H (ICRP) | M (Brown) | H (Peters) | R (Peters) | H (Shah) | M (Shah) | R (Shah) |
|---|---|---|---|---|---|---|---|
| stomach | 0.144 [12] | 0.000219 [2] | 0.154 [3] | 0.0011 [3] | 0.144 [12] | 0.00219 [2] | 0.0011 [3] |
| smallIntestine | 0.625 [12] | 0.000327 [2] | 1.652 [3] | 0.01 [3] | 0.385 [4] | 0.000728 [4] | 0.00499 [4] |
| largeIntestine | 0.356 [12] | 0.000183 [2] | | | 0.548 [4] | 0.000314 [4] | 0.00287 [4] |
| pancreas | 0.133 [12] | 0.000299 [2] | 0.084 [3] | 0.0013 [3] | 0.104 [4] | 0.000097 [4] | 0.001 [4] |
| spleen | 0.144 [12] | 0.0000857 [2] | 0.189 [3] | 0.0006 [3] | 0.221 [4] | 0.000127 [4] | 0.00277 [4] |
| liver | 1.714 [12] | 0.00135 [2] | 1.687 [3] | 0.0103 [3] | 2.143 [4] | 0.00193 [4] | 0.0157 [4] |
| brain | 1.381 [12] | 0.000417 [2] | 1.449 [3] | 0.0017 [3] | 1.45 [4] | 0.000485 [4] | 0.00228 [4] |
| heart | 0.314 [12] | 0.000123 [2] | 0.266 [3] | 0.0008 [3] | 0.341 [4] | 0.000152 [4] | 0.00102 [4] |
| lungs | 1.142 [1] | 0.000172 [2] | 1.169 [3] | 0.001 [3] | 1 [4] | 0.000204 [4] | 0.0014 [4] |
| kidneys | 0.295 [12] | 0.000417 [2] | 0.308 [3] | 0.0023 [3] | 0.332 [4] | 0.000525 [4] | 0.00241 [4] |
| fat | 19.791 [9, 13] | 0.00172 [2] | 10.01 [3] | 0.01 [3] | 13.465 [4] | 0.00198 [4] | 0.0331 [4] |
| skin | 3.562 [12] | 0.00404 [2] | 7.77 [3] | 0.04 [3] | 3.408 [4] | 0.00502 [4] | 0.0499 [4] |
| skeletalMuscle | 27.619 [12] | 0.00941 [2] | 30.03 [3] | 0.122 [3] | 30.078 [4] | 0.0113 [4] | 0.122 [4] |
| skeleton | 7.726 [12] | 0.00262 [2] | 8.68 [3] | 0.0158 [3] | 10.165 [4] | 0.00282 [4] | 0.021 [4] |
| redMarrow | 1.138 [5] | 0.000375 [10] | | | 1.138 [5] | 0.000375 [10] | 0.0037 [10] |
| thyroid | 0.0192 [12] | | | | | | |
| gonads | 0.0168 [12] | | 0.0357 [3] | 0.0025 [3] | | | |
| adrenals | 0.0168 [12] | | | | | | |
| bladder | 0.0481 [12] | | | | | | |
| lymphNodes | 0.709 [12] | 0.00174 [14, 15] | 0.709 [12] | 0.00115 [4] | 0.274 [4] | 0.000113 [4] | 0.00115 [4] |
| thymus | | | | | 0.00641 [4] | 0.000009 [4] | 0.000096 [4] |
| misc | 2.859 [12] | 0.0174 [2, 6, 7] | | | 4.852 [4] | 0.000465 [4] | 0.00609 [4] |

**Table S4.** Organ volume in litres or $dm^3$ (H: Human, M: Mouse, R: Rat).

The absolute difference in delivery rates could be quite large, dependent on the organ. For example, absolute lung delivery rates in the rat are predicted to be approximately 11 times higher than in humans if the same blood concentration of T-cells is assumed, or 2400 times higher if the same number of cells is introduced for each species. If the same concentration or number of cells is introduced to a mouse and a human, the lung delivery rate per $mm^3$ would be 8.4 or 17700 times higher in mice. The absolute rates, assuming the same number of CART-cells ($10^8$) are introduced, are shown in main text table 1. As before, the flow from both the hepatic artery and portal vein are included in the calculated delivery rates per $mm^3$ to the liver, and the pulmonary circuit and lung blood supply are are both included in the calculated rates for the lung.

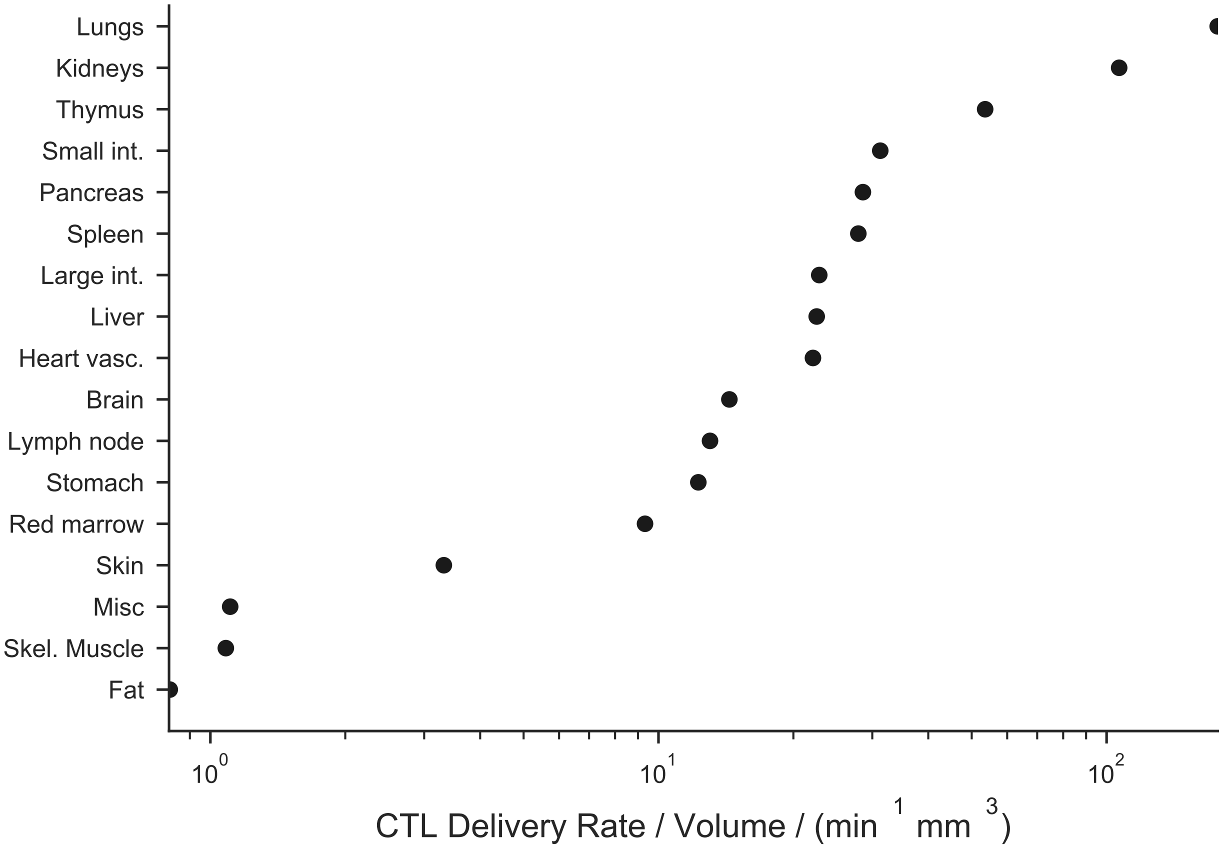


**Figure S1. Maximal CART-cell delivery rates vary by organ:** CART-cell delivery rates per volume (cells/min/$mm^3$) to human organs were computed by the model, and are approximately equal to the product of organ perfusion and assumed blood CART-cell concentration. T-cell extravasation rates cannot exceed their rate of delivery by the blood; the maximum extravasation rates would be obtained in the case that all T-cells entering the organ vasculature extravasate. Note that the lungs and kidneys have the highest rates of T-cell delivery.

#### B.3 Predicted delivery rates of other immune populations in humans

The results of the model are relevant not just to CART-cells but to any immune cell population of a known blood concentration or count. The data in main text table 1 may be used by multiplying results by total blood volume and the blood concentration, or count, of an immune cell population of interest. The mean and range of observed values [16, 17] for several immune cell blood concentrations are shown in supplementary table S5. For example, assuming a human CTL concentration of $C = 3 \times 10^3$ cells/μL [16, 17] and a total blood volume of 5.3L [1] yields a total number of cells equal to approximately $1.6 \times 10^{10}$ cells, or 160 times the values quoted for CART-cells in this study. This results in maximum delivery rates 160 times larger than those presented in the main text. This, however, assumes that *all* blood CTLs are relevant to the target of interest.

| Cell Type | Rate Relative to CARs | Biological Variation |
|---|---|---|
| B Cells | 240× | 24× – 400× |
| Total T Cells | 400× | 64× – 560× |
| CD4+ T cells (Th) | 350× | 130× – 540× |
| CD8+ T cells (CTL) | 160× | 32× – 410× |
| NK cells | 95× | 0× – 350× |
| Monocytes | 80× | 0× – 290× |

**Table S5. Relative delivery rates in humans by immune cell type:** Delivery rates of immune cells to organs are proportional to blood immune cell concentrations, thus, predicted delivery rates scale trivially when a different cell concentration is considered. Published data on human blood immune cell concentrations [16, 17] were used to calculate mean delivery rates and inter-individual variation for B, T helper, cytotoxic T-lymphocyte (CTL), natural killer and monocyte cells, relative to the values given for CART-cells in the main text.

#### B.4 Use of different parameter sources

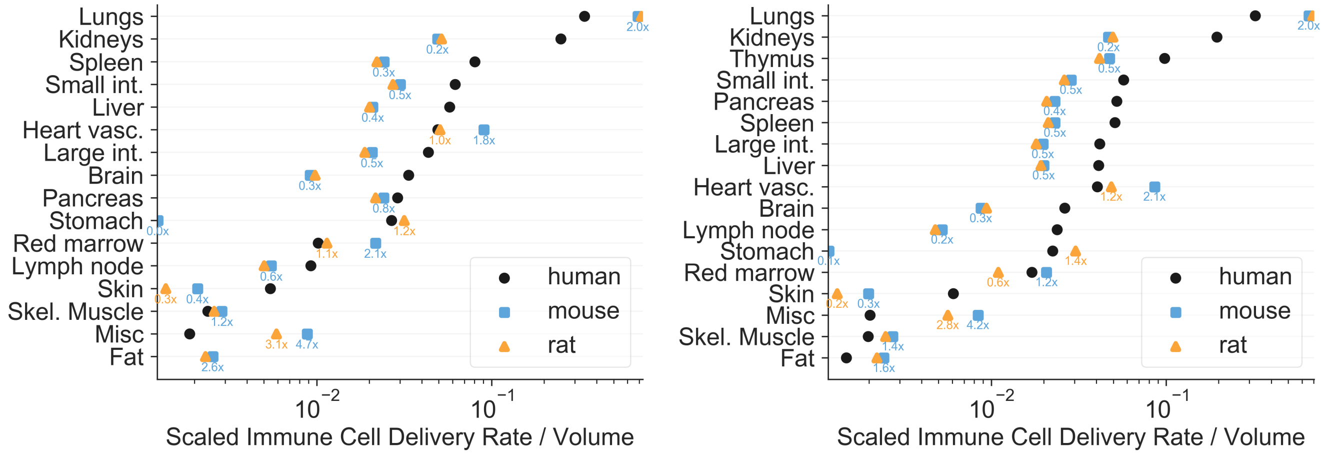


**Figure S2. Relative predicted delivery rates to non-tumour tissue in organs in humans, rats and mice.** Rates within each species were normalised to sum to 1.0 to give relative values for comparison, as in main text figure 2. Numerical values (text labels) give the mean predicted delivery rates in rats and mice relative to human values, with colours matching the legend. The left plot uses human predictions parameterised from the ICRP anatomical reference [1], and the right plot uses data from Shah *et al*'s compilation [4]. Note the different ordering of organs, and that the horizontal axis is a log scale.

As discussed in the main text, the conclusions of an anatomical model such as this depend strongly on the values of parameters used. This point may be seen more clearly if two different highly cited anatomical references are plotted side by side. Supplementary figure S2 shows the same predictions for the mouse and human, but the left plot uses the ICRP's highly cited anatomical reference for human parameters [1] and the right plot uses the compilation by Shah *et al* [4]. Note that both the ordering and the scaling of data is slightly different.

#### B.5 Tumour perfusion compared to healthy tissue

To quantify the delivery rate to tumours, per volume of tumour (cells/min/mm$^3$), the perfusion of different tumour types was required. The data sourced from literature [18–63] are presented in supplementary figure S3. Tumour perfusion can be greater or lower than that of the source tissue, varying by many orders of magnitude. The range is likely to represent tumours that do or do not secrete angiogenic factors, or whose new vasculature has been renormalised or not. As discussed in the main text, whether tumour perfusion typically outstrips that of the bearing organ appears to be linked to the function of the organ itself, *e.g.* the kidneys have evolved to have very efficient perfusion that is unlikely to be outstripped by random, disordered angiogenesis.

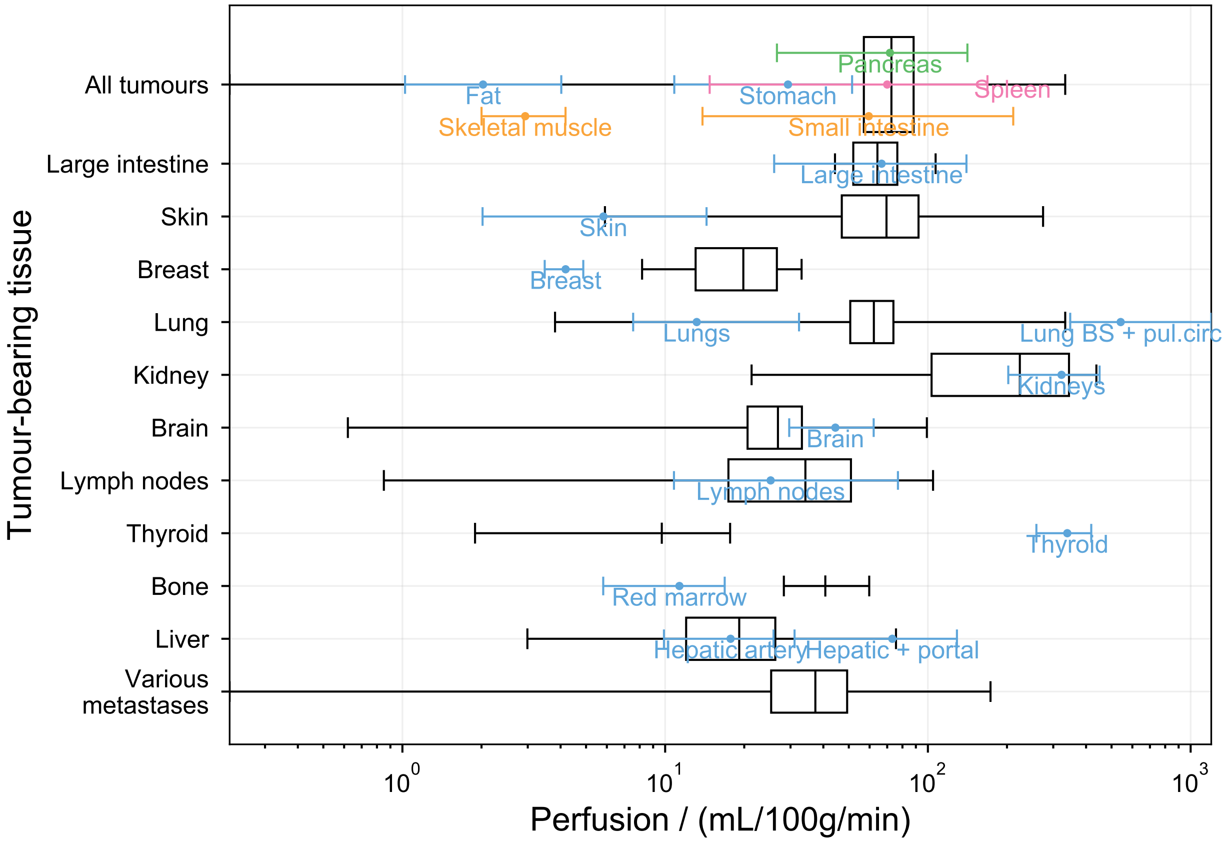


**Figure S3. Perfusion of human tumours:** Tumour perfusion sourced from literature [18–63]. As in main text figure 3, data are arranged by tumour location, with all lung-located tumours in the 'Lung' column. Black boxes represent the mean ± standard deviation of tumour perfusion and whiskers indicate the range of tumour perfusion found in the literature. Coloured error bars represent the mean and range of perfusion of healthy tissue from the data used in main text figure 2. Note that the horizontal axis is logarithmic. "Lung BS + pul.circ" indicates the perfusion of the lung when blood from both the lung blood supply and the pulmonary are considered, whereas the "Lungs" entry indicates the perfusion when only the blood supply is considered.

#### B.6 Sensitivity analysis

The reported results can be obtained from an ODE model of the circulatory system, described in supplementary section A.1. To do this, blood concentration is assumed to be constant. To ensure that this is true, extravasation probabilities for all organs are set to zero but for a unit volume in an organ of interest. To show that setting other values to zero does not affect results, we performed a sensitivity analysis on the model. Random values of the extravasation probabilities of T-cells into all 20 organs and the "tumour" ($1mm^3$ volume of interest) between 9% and 11% are chosen. Setting the tumour to be in each organ in turn, the extravasation rate of T-cells into the tumour is calculated for all organs. This process was repeated over 1000 simulations and the data was used to grow a random forest for each organ, using Scikit [64]. This process produces indices between 0 and 1 which indicate the 'importance' of each parameter for prediction of the extravasation rate of T-cells into a tumour in a given organ. On average, the importance factor of the tumour-specific extravasation proportion was 0.91. The importance factor of all other organs, including the tumour-bearing organ, was 0.004. The average pearson rank correlation between the extravasation rate of T-cells into the tumour and the tumour-specific extravasation proportion was 0.95, and -0.04 with the extravasation proportion into all other organs. Exact data are reproduced in supplementary table S6.

| Extrvstn prob into↓ | Tumour bearing organ (predicted delivery rate to organ): Sto | SI | LI | Pan | Spl | Liv | Bra | Hrt | Lng | Kid | Fat | Skn | Mus | Bon | Thy | Gnd | Adr | Bla | LNs | Mis |
|---|---|---|---|---|---|---|---|---|---|---|---|---|---|---|---|---|---|---|---|---|
| Tumour | 0.916 | 0.909 | 0.911 | 0.912 | 0.909 | 0.907 | 0.905 | 0.908 | 0.906 | 0.906 | 0.916 | 0.911 | 0.914 | 0.919 | 0.906 | 0.913 | 0.906 | 0.908 | 0.905 | 0.919 |
| Stomach | 0.001 | 0.001 | 0.001 | 0.001 | 0.001 | 0.001 | 0.001 | 0.001 | 0.001 | 0.001 | 0.001 | 0.001 | 0.001 | 0.001 | 0.001 | 0.001 | 0.001 | 0.001 | 0.001 | 0.001 |
| Small intestine | 0.001 | 0.001 | 0.001 | 0.001 | 0.001 | 0.003 | 0.001 | 0.001 | 0.001 | 0.001 | 0.001 | 0.001 | 0.001 | 0.001 | 0.001 | 0.001 | 0.001 | 0.001 | 0.001 | 0.001 |
| Large intestine | 0.001 | 0.001 | 0.001 | 0.001 | 0.001 | 0.002 | 0.001 | 0.001 | 0.001 | 0.001 | 0.001 | 0.001 | 0.001 | 0.001 | 0.001 | 0.001 | 0.001 | 0.001 | 0.001 | 0.001 |
| Pancreas | 0.001 | 0.001 | 0.001 | 0.001 | 0.001 | 0.001 | 0.001 | 0.001 | 0.001 | 0.001 | 0.001 | 0.001 | 0.001 | 0.001 | 0.001 | 0.001 | 0.001 | 0.001 | 0.001 | 0.001 |
| Spleen | 0.001 | 0.001 | 0.001 | 0.001 | 0.001 | 0.001 | 0.001 | 0.001 | 0.001 | 0.001 | 0.001 | 0.001 | 0.001 | 0.001 | 0.001 | 0.001 | 0.001 | 0.001 | 0.001 | 0.001 |
| Liver | 0.004 | 0.004 | 0.004 | 0.004 | 0.004 | 0.005 | 0.005 | 0.004 | 0.005 | 0.004 | 0.004 | 0.005 | 0.004 | 0.004 | 0.004 | 0.004 | 0.005 | 0.005 | 0.005 | 0.004 |
| Brain | 0.002 | 0.002 | 0.002 | 0.002 | 0.002 | 0.002 | 0.002 | 0.002 | 0.002 | 0.002 | 0.002 | 0.002 | 0.002 | 0.002 | 0.002 | 0.002 | 0.002 | 0.002 | 0.002 | 0.002 |
| Heart vasculature | 0.001 | 0.001 | 0.001 | 0.001 | 0.001 | 0.001 | 0.001 | 0.001 | 0.001 | 0.001 | 0.001 | 0.001 | 0.001 | 0.001 | 0.001 | 0.001 | 0.001 | 0.001 | 0.001 | 0.001 |
| Pulmonary tissue | 0.003 | 0.003 | 0.003 | 0.003 | 0.002 | 0.002 | 0.003 | 0.003 | 0.003 | 0.003 | 0.002 | 0.002 | 0.003 | 0.002 | 0.003 | 0.002 | 0.003 | 0.003 | 0.003 | 0.002 |
| Kidneys | 0.001 | 0.001 | 0.001 | 0.001 | 0.001 | 0.001 | 0.001 | 0.001 | 0.001 | 0.001 | 0.001 | 0.001 | 0.001 | 0.001 | 0.001 | 0.001 | 0.001 | 0.001 | 0.001 | 0.001 |
| Fat | 0.004 | 0.004 | 0.004 | 0.004 | 0.004 | 0.004 | 0.004 | 0.004 | 0.004 | 0.004 | 0.004 | 0.004 | 0.004 | 0.003 | 0.004 | 0.004 | 0.004 | 0.004 | 0.004 | 0.003 |
| Skin | 0.002 | 0.003 | 0.002 | 0.002 | 0.002 | 0.003 | 0.003 | 0.003 | 0.003 | 0.003 | 0.002 | 0.003 | 0.003 | 0.002 | 0.002 | 0.002 | 0.003 | 0.003 | 0.002 | 0.002 |
| Skeletal muscle | 0.053 | 0.058 | 0.057 | 0.056 | 0.058 | 0.055 | 0.061 | 0.058 | 0.060 | 0.060 | 0.053 | 0.057 | 0.055 | 0.051 | 0.061 | 0.055 | 0.060 | 0.058 | 0.061 | 0.052 |
| Skeleton | 0.003 | 0.003 | 0.003 | 0.003 | 0.003 | 0.004 | 0.003 | 0.003 | 0.004 | 0.003 | 0.003 | 0.003 | 0.003 | 0.003 | 0.004 | 0.003 | 0.003 | 0.003 | 0.004 | 0.003 |
| Thyroid | 0.001 | 0.001 | 0.001 | 0.001 | 0.001 | 0.001 | 0.001 | 0.001 | 0.001 | 0.001 | 0.001 | 0.001 | 0.001 | 0.001 | 0.001 | 0.001 | 0.001 | 0.001 | 0.001 | 0.001 |
| Gonads | 0.001 | 0.001 | 0.001 | 0.001 | 0.001 | 0.001 | 0.001 | 0.001 | 0.001 | 0.001 | 0.001 | 0.001 | 0.001 | 0.001 | 0.001 | 0.001 | 0.001 | 0.001 | 0.001 | 0.001 |
| Adrenals | 0.001 | 0.001 | 0.001 | 0.001 | 0.001 | 0.001 | 0.001 | 0.001 | 0.001 | 0.001 | 0.001 | 0.001 | 0.001 | 0.001 | 0.001 | 0.001 | 0.001 | 0.001 | 0.001 | 0.001 |
| Urinary bladder | 0.001 | 0.001 | 0.001 | 0.001 | 0.001 | 0.001 | 0.001 | 0.001 | 0.001 | 0.002 | 0.001 | 0.001 | 0.001 | 0.001 | 0.001 | 0.001 | 0.001 | 0.001 | 0.001 | 0.001 |
| Lymph node | 0.001 | 0.001 | 0.001 | 0.001 | 0.001 | 0.001 | 0.001 | 0.001 | 0.001 | 0.001 | 0.001 | 0.001 | 0.001 | 0.001 | 0.001 | 0.001 | 0.001 | 0.001 | 0.001 | 0.001 |
| Misc | 0.001 | 0.001 | 0.001 | 0.001 | 0.001 | 0.001 | 0.001 | 0.001 | 0.001 | 0.001 | 0.001 | 0.001 | 0.001 | 0.001 | 0.001 | 0.001 | 0.001 | 0.001 | 0.001 | 0.001 |

**Table S6.** Importance indices derived from the random forest algorithm (see text) for each the extravasation proportion of T-cells into each organ, for each tumour extravasation rate. 0 means no effect on the observable and 1 is maximal importance. Columns are abbreviations of the organs as written on the left of each row, starting from “Sto” for “Stomach”.
